# Acquisition of novel arrays via horizontal gene transfer rewire CRISPR-mediated defense in *Pseudomonas aeruginosa*

**DOI:** 10.64898/2026.04.27.721218

**Authors:** Robbie Paul P. Malaluan, Ron Leonard V. Dy

## Abstract

CRISPR-Cas systems form the adaptive immunity of prokaryotes, conferring sequence-specific protection against genetic parasites. Here, we functionally characterized the CRISPR-Cas system of *Pseudomonas aeruginosa* ATCC 10145 (PA10145), which led us to discover the existence of an isolated CRISPR array, unique to this system. PA10145 possesses a type I-F CRISPR-Cas composed of a *cas* operon flanked by two divergently organized CRISPRs. The isolated CRISPR array, CRISPR3, is located ∼1.3 million bp away from the *cas* loci. The *cas* and three CRISPR arrays are active. Plasmids with an engineered protospacer matching any of the three arrays were targeted and stimulated hyperactive adaptation in all CRISPR arrays of PA10145 if the plasmids possessed an intact protospacer adjacent motif (PAM), whereas minimal to no adaptation was observed when PAM was mutated. Spacer acquisition via interference-driven adaptation proceeds through strand-biased priming in PA10145. Interestingly, the isolated CRISPR3 and the *cas*-adjacent CRISPR2 have nearly identical leader sequences with only 3 bp mismatches. From a survey of CRISPR loci in 1,198 *P. aeruginosa* genomes, isolated arrays only occur as type I-F with similarly matching leaders to CRISPR2. Highly-transmissible mobile genetic elements (MGEs) associate with CRISPR2 and CRISPR3, suggesting that isolated arrays might have originated from recombination events involving CRISPR2 as facilitated by these MGEs. Tracing evolutionary trajectories of the isolated CRISPR3 relative to *cas*-adjacent arrays revealed that CRISPR3 is laterally transferred across *P. aeruginosa* genomes. Taken together, these results implicate the role of horizontally-acquired isolated arrays in CRISPR-mediated pan-immunity as gateways to mobilize genetic memories.

## Introduction

Clustered Regularly Interspaced Short Palindromic Repeats (CRISPRs) and its CRISPR-associated (*cas*) genes constitute the adaptive immune system in bacteria and Archaea [1]. These systems are typically composed of a CRISPR array, consisting of short repeats separated by spacers derived from invading mobile genetic elements (MGEs) such as plasmids and bacteriophages, and adjacent *cas* genes organized in one or more operons. To impart immunity, CRISPR-Cas functions in three sequential stages [1]. During adaptation, short fragments of the invading genetic material are integrated as new spacers into the CRISPR array, thereby updating the genetic memory of immunity. The CRISPR array is then transcribed [2,3] and processed by specific endonucleases into mature CRISPR RNAs (crRNAs), each consisting of a spacer sequence flanked by remnants of the repeats [4]. Finally, Cas proteins form ribonucleoprotein complexes with mature crRNAs, which guide the interference machinery to recognize and degrade complementary foreign genetic materials, effectively preventing their proliferation [4]. CRISPR-Cas systems are highly abundant in bacterial genomes and are currently classified into two major Classes (Class 1 and Class 2), which are further subdivided into seven distinct types (Type I to VII) and over 30 subtypes based on their signature Cas proteins and mechanistic differences [5,6]. The detailed characterization of these systems, along with their natural diversity, enabled the development of revolutionary genome editing and targeting tools with far-reaching biotechnological applications.

Beyond their roles in immunity, CRISPR-Cas systems contribute significantly to bacterial evolution by driving genome rearrangements [7], shaping microbial population dynamics [8–10], and balancing horizontal gene transfer [11–13]. One apparent paradox is that while CRISPR-Cas protect their hosts by restricting all three routes of HGT, namely transformation [14,15], conjugation [16–18] and transduction [11,19]; bacteria themselves rely on HGT for their rapid evolutionary adaptation, such as the acquisition of virulence genes and the diversification of CRISPR-Cas. This conflict suggests that evolutionary trade-offs may shape the maintenance and dissemination of CRISPR-Cas systems. Indeed, their distribution may be influenced by ecological factors that alter the selective pressure for CRISPR-mediated immunity, such as populational density [20], viral load [21,22], and the structure of microbial communities [8–10]. Hence, the diverse CRISPR-Cas architectures we observe today likely reflect the combined effects of these interrelated drivers, which led to the emergence of CRISPR-Cas variants that lack major components, such as the absence of functional interference modules, CRISPR arrays, or entire *cas* operons.

CRISPR arrays are typically located adjacent to *cas* genes, reflecting their close functional relationship and co-evolution [1]. However, a substantial subset of CRISPR arrays is found without neighboring *cas* genes [23,24]. These are classified as either orphaned, found in genomes entirely lacking *cas* genes, or isolated, which are arrays located at genomic loci distant from *cas* operons. Orphaned and isolated CRISPR arrays are thought to originate from three proposed mechanisms: MGE-mediated transfer, excision of the *cas* operon leaving behind the CRISPR, or off-target spacer acquisition events [24]. Despite being viewed as evolutionary vestiges, some of these standalone CRISPR arrays might remain functional. For example, isolated CRISPR arrays, although separated from their cognate *cas* genes, should still be able to participate in immunity [25,26]. Meanwhile, in *Escherichia coli*, an orphan CRISPR array containing spacers targeting its native *cas* genes was shown to inhibit acquisition of the type I-F CRISPR-Cas system, thereby preventing uptake of foreign *cas* operons [27]. Understanding the origins of these isolated arrays is critical because they could offer insights into the evolution of CRISPR-Cas systems and their potential for novel forms of CRISPR-based regulations.

Owing to the high genomic plasticity of *Pseudomonas aeruginosa* (*Psa*), which features an abundance of MGEs [28,29], the opportunistic human pathogen could be a suitable model to investigate the interplay between MGEs and CRISPR-Cas. The *Psa* strain ATCC 10145 (DSM 50071), abbreviated in this study as PA10145, contains a type I-F CRISPR-Cas system with the characteristic genetic architecture similar to the model PA14 strain [30–37]. Its type I-F CRISPR-Cas loci consists of a *cas* operon flanked by two divergently organized CRISPR arrays. With respect to the transcriptional direction of the *cas* operon, the upstream array is designated as CRISPR2 (CR2) while the downstream locus is CRISPR1 (CR1) [30]. Notably, PA10145 harbors an isolated CRISPR array, hereafter referred to as CRISPR3 (CR3), located approximately 1.3 Mbp away from the operon (**Figure 1**). The emergence and involvement of an isolated CRISPR array situated millions of bp away from its cognate CRISPR-Cas in *Psa* remains to be understood. Thus, in this study, the functionality of the isolated CR3 towards adaptive immunity was investigated. Specifically, we demonstrate the involvement of CR3 in imparting MGE-interference and inducing rapid acquisition of new spacers via primed adaptation. We also present evidence for the dissemination of isolated arrays in *Psa* via horizontal gene transfer through comparative genomics of more than 1,000 published *Psa* genomes.

**Figure 1.**
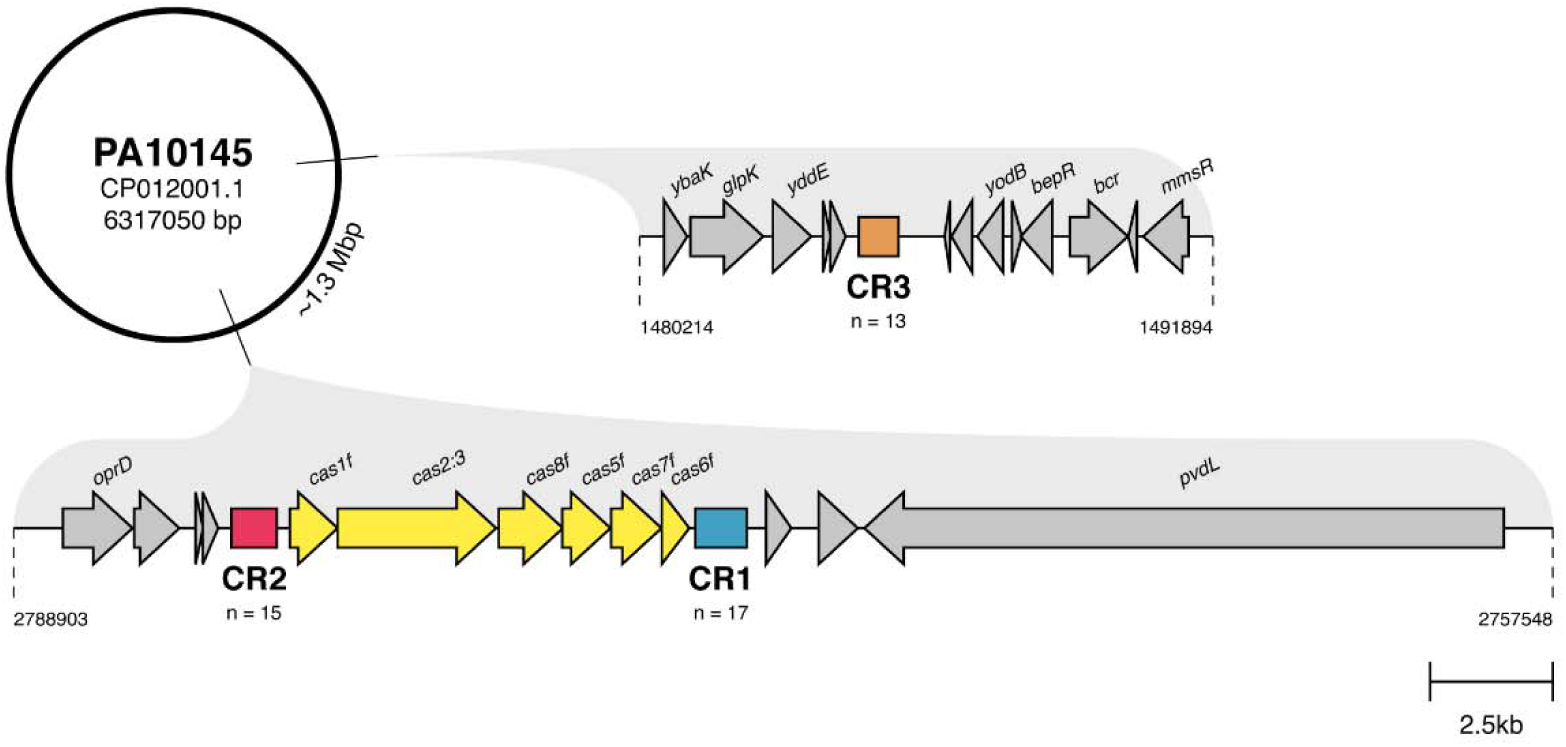
The CRISPR-Cas system of PA10145. The type I-F CRISPR-Cas system of PA10145 is located between 2773971 and 2784471 under the NCBI GenBank Accession No. CP012001.1. Two adjacent CRISPR arrays flank the *cas* operon. A distal and isolated CR3 array is located ∼1.3 Mbp away from the *cas* operon.

## Results

### An isolated array functions in CRISPR-Cas immunity

The isolated CR3 array of PA10145 contains 13 unique spacers interspaced by a consensus repeat sequence identical to those of the adjacent CR1 and CR2 repeats, except for a 1 base-pair mismatch at the predicted loop region (**Figure S1**). The adjacent CR2 and isolated CR3 share 98.01% sequence identity across an upstream 900 bp region (**Figure S2**). This includes their presumed leader sequences which were nearly identical, differing at only three base-pair positions (**Figure S3**), yet the spacers they harbor are entirely distinct (**Figure S4**). All three CRISPR arrays are predicted to target foreign genetic elements such as phages and plasmids (**Table S1-S3**) [38]. The strong conservation of these key genetic features across the isolated CR3 and the adjacent CRISPR arrays suggests that all three are functionally linked and form the CRISPR-Cas system of PA10145.

We then examined the role of CR3 across the three aspects of CRISPR-Cas immunity: biogenesis, interference, and adaptation. To assess expression of type I-F CRISPR-Cas natively in PA10145, the upstream intergenic regions (UIRs) of each component were fused to a plasmid-borne promoterless *lacZ* reporter (**Figure S5A**). Fluorometric MUG assays revealed that the UIRs of *cas,* CR2, CR1, and CR3 all drove *lacZ* expression, indicating that each component harbors an active promoter (**Figure S5B**). Notably, CR2 and CR3 exhibited comparable and higher promoter activities relative to *cas* and CR1, consistent with their nearly identical UIRs (**Figure S5B**). Collectively, these results indicate that the isolated CR3 array is expressed at levels similar to the core type I-F CRISPR-Cas components in PA10145.

To determine the participation of the isolated CR3 to CRISPR-Cas mediated interference of MGEs, we engineered plasmids each carrying a single protospacer (PS) derived from one of the three CRISPR arrays of PA10145 [39]. Targeting (T) plasmids were designed to contain a PS flanked by the functional 3’-GG PAM [35,36], while non-targeting (NT) variants carried a mutated 3’-GT PAM [39]. As expected, T plasmids carrying PS sequences recognized by CR2 or CR1 had up to ∼50% reduction in transformation efficiency, while their NT counterparts transformed at efficiencies comparable to the no PS (-) control (**Figure 2A**). The T-CR3 plasmid was similarly targeted, exhibiting a ∼68% reduction in transformation efficiency (**Figure 2A**). PAM mutation in the NT-CR3 plasmid attenuated targeting; however, some interference relative to the no PS plasmid control was still observed (**Figure 2A**). These results demonstrate that all three CRISPR arrays of PA10145, including the isolated CR3, are functional and capable of mediating plasmid interference.

**Figure 2.**
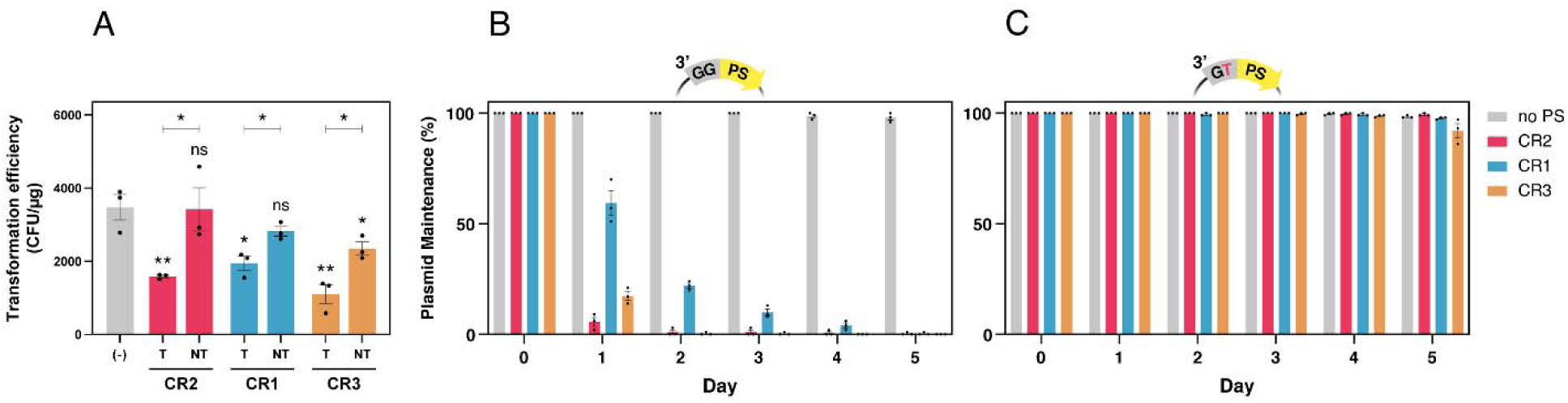
The isolated CR3 participates in type I-F CRISPR-Cas immunity of PA10145. (A) Transformants per 1 μg of plasmid were calculated in triplicate for each plasmid construct. When flanked with an intact 3’-GG PAM, a single PS recognized by CR2, CR1, and CR3 significantly decreased the transformation efficiency of the T plasmids (p = 0.006, p=0.019, and p=0.006 respectively) relative to the no PS (-) control. When PAM was mutated, transformation efficiencies of NT-CR2, NT-CR1, and NT-CR3 were restored compared to their corresponding T vectors (p=0.036, p=0.021, and p=0.018 respectively). Notably, only the NT-CR3 vector resulted in a significant reduction in transformation efficiency relative to no PS (p=0.046). (B-C) Percent plasmid maintenance per day was determined by the number of surviving colonies on the S plate. (B) T plasmids were significantly degraded by day 1. T-CR2 or T-CR3 were rapidly lost while a gradual decrease can be seen for T-CR1. (C) The non-canonical 3’-GT PAM in all NT constructs restored maintenance across 5 days. (A) two-tailed Student’s t-test; **p < 0.01; *p < 0.05. Significance directly above experimental bars indicates comparison to the no PS (-) control (A-D) Bars represent mean ± SEM.

Primed adaptation enables rapid spacer acquisition following initial target recognition [39]. To determine whether CR3-mediated recognition can trigger adaptation, we performed a plasmid loss assay in which bacteria carrying T or NT plasmids were passaged for five days without antibiotic selection. T-CR2 and T-CR3 plasmids were rapidly lost in the bacterial population within 2 days, whereas the T-CR1 plasmid was lost more gradually, disappearing by day 5 (**Figure 2B**). In contrast, all NT plasmids were stably maintained throughout the 5-day timeframe (**Figure 2C**). To confirm that plasmid loss was accompanied by CRISPR adaptation, we PCR-screened colonies that had lost the plasmid on day 5 for expansion across the three CRISPR arrays. Spacer acquisition was detected in approximately 55% (5/9), 88% (8/9), 44% (4/9) of colonies that lost T-CR2, T-CR1, and T-CR3 plasmids, respectively (**Figure S6A-C**). Notably, colony 2 from the T-CR1 set-up acquired seven new spacers distributed across all three CRISPR arrays (**Figure S6B)**. Among the NT plasmids, colonies that did lose the plasmid also showed evidence of CRISPR expansion: 33% (3/9) and 66% (6/9) of colonies from the NT-CR1 and NT-CR3 set-ups, respectively, had acquired new spacers, while no adaptation was detected in NT-CR2 (**Figure S6D-F**). These findings demonstrate that regardless of which array mediated target recognition, plasmid loss can trigger spacer acquisition across all three CRISPR arrays in PA10145.

### Interference drives primed adaptation in the type I-F CRISPR-Cas of PA10145

To identify the main modality for rapid spacer acquisition in the type I-F CRISPR-Cas system of PA10145, we screened bacterial populations on day 5 post-exposure to the T or NT plasmids for CRISPR array expansion. Strikingly, expansion across all three arrays was most pronounced when PA10145 was challenged with T plasmids, regardless of which array targeted the PS (**Figure 3A**), whereas minimal to no adaptation occurred in their NT counterparts (**Figure 3B**). Prolonged incubation with the no PS vector similarly failed to induce naïve adaptation (**Figure S7**). Together, these results establish that interference drives primed adaptation in the type I-F CRISPR-Cas of PA10145.

**Figure 3.**
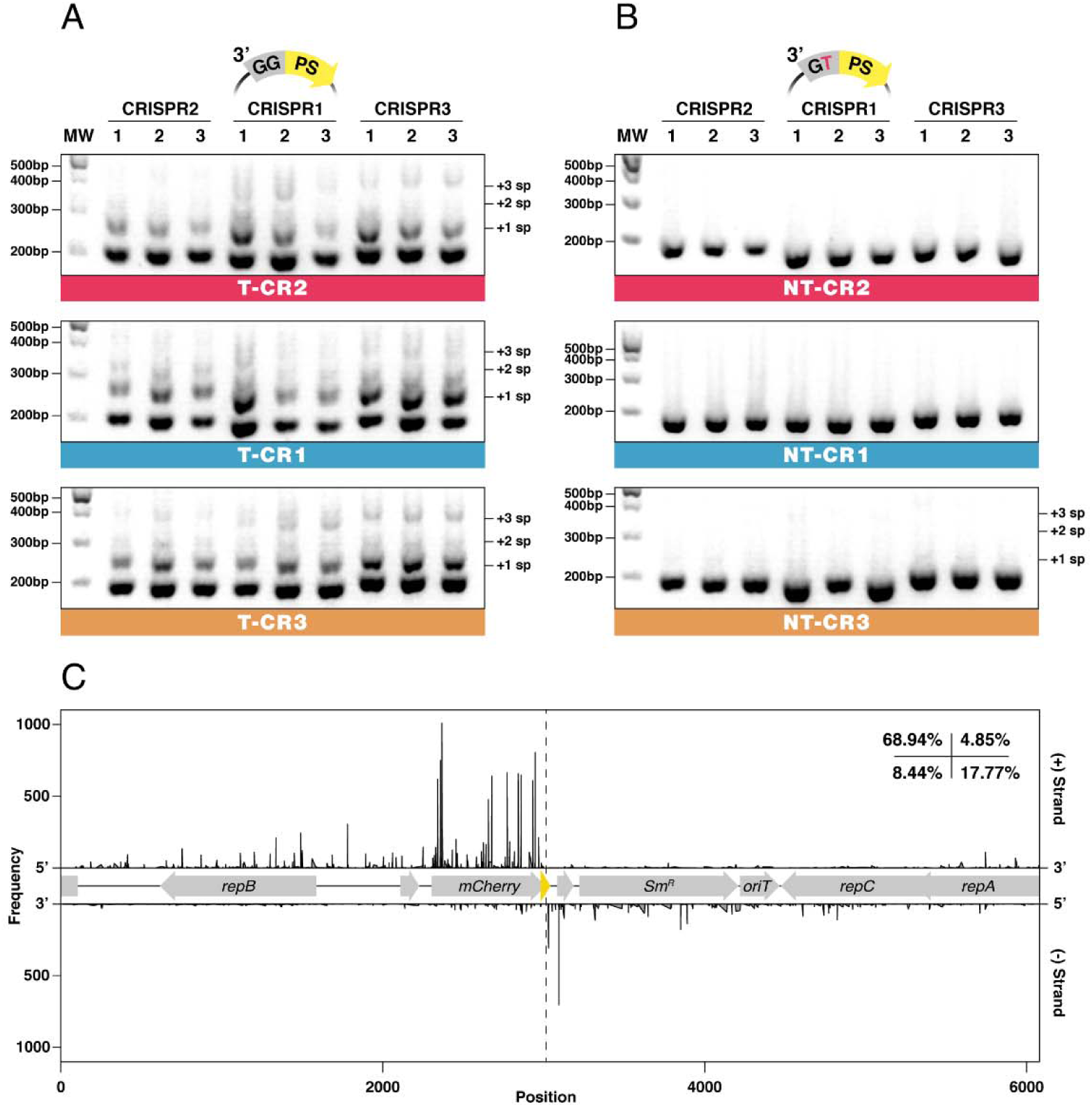
Interference drives rapid spacer acquisition through strand-biased priming. (A-B) PCR templates were sampled from the bacterial culture at day 5 post-exposure to the PS-containing plasmids per biological replicate. (A) CRISPR expansion was most visible in AGE profiles of all populations exposed to T plasmids. (B) Laddering could not be detected when PAM was mutated in NT-CR2 and NT-CR1 while faint heavier bands were observed in NT-CR3. (C) Sequencing the CRISPR expansions from all T set-ups revealed distributions of novel spacers upon mapping to the T plasmid. The targeted protospacer is located on the (-) strand in all set-ups; therefore, it follows that the target strand is the (-) strand while the non-target strand is the (+) strand. Percentages on the top right correspond to the portion of spacers mapped to the non-target (+) and target (-) strand, and the 5’ and 3’ direction with respect to the target protospacer marked by the center dashed line.

To further characterize the mechanism underlying primed adaptation following CRISPR-mediated plasmid clearance, we sequenced the PCR amplicons from expanded CRISPR arrays. Analysis of over ∼200,000 sequenced amplicons revealed that the extent of expansion was consistent across arrays, with the vast majority (∼95%) having acquired only a single new spacer (**Figure S8**). Newly acquired spacers were distributed evenly across the adjacent CR1 and CR2 arrays and the isolated CR3 (**Figure S8, Table 1**). In total, 25,112 newly-acquired spacers mapped to the T plasmid, with none matching the PA10145 genome. These spacers predominantly conformed to the canonical 32 bp length and mapped to protospacers flanked by a 3’-GG PAM (**Figure S9**). The decline in acquisitional frequency in the 5’ direction on both strands, relative to the original PS target, is consistent with the bidirectional modality of interference-driven primed adaptation [39,40] (**Figure 3C**). Since the original PS targets were designed to be on the target (-) strand, the predominant mapping of novel spacers (∼75%) to the (+) strand indicates a preferential acquisitional bias for the non-target strand during acquisition (**Table 1**). Importantly, spacer distributions were consistent across all three arrays, confirming that the isolated CR3 actively participates in interference-mediated primed adaptation alongside CR1 and CR2 (**Figure S10**).

**Table 1.**
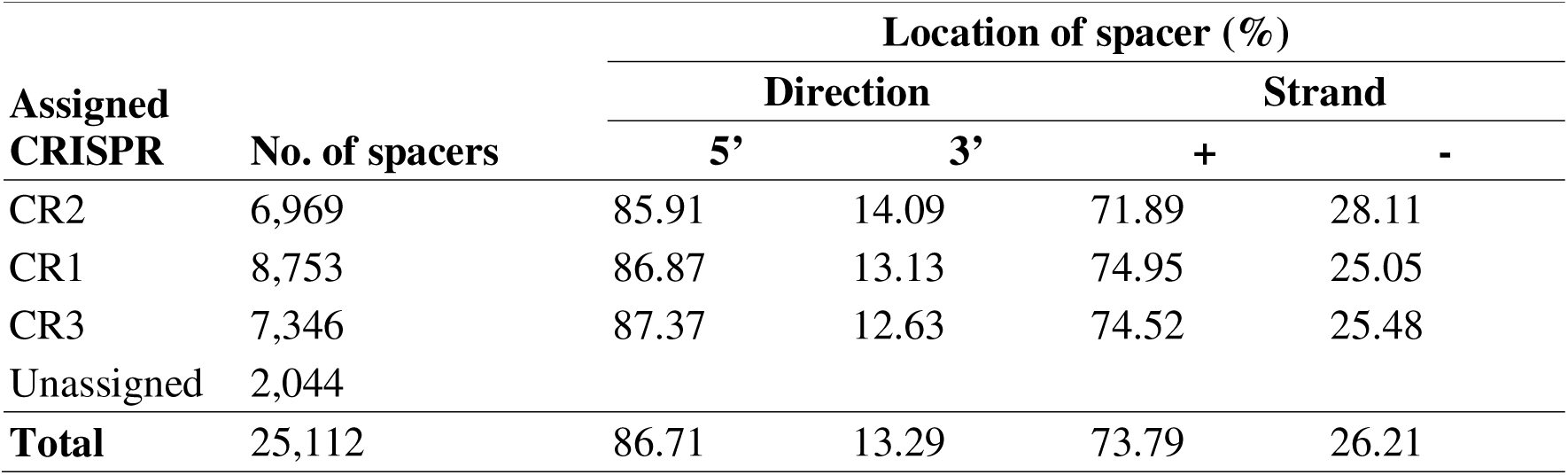
Distribution of novel spacers relative to the target protospacer. Unassigned spacers were mappable to the targeted vector but could not be definitively binned to a CRISPR due to sequencing errors in the two pre-existing spacers from its read.

| Assigned<br>CRISPR | No. of spacers | Location of spacer (%) |  |  |  |
| --- | --- | --- | --- | --- | --- |
|  |  | Direction |  | Strand |  |
|  |  | 5' | 3' | + | - |
| CR2 | 6,969 | 85.91 | 14.09 | 71.89 | 28.11 |
| CR1 | 8,753 | 86.87 | 13.13 | 74.95 | 25.05 |
| CR3 | 7,346 | 87.37 | 12.63 | 74.52 | 25.48 |
| Unassigned | 2,044 |  |  |  |  |
| **Total** | 25,112 | 86.71 | 13.29 | 73.79 | 26.21 |

To gain deeper insights into the sequential dynamics of spacer acquisition during priming, we hypothesized that the second acquired spacer (+2) would exhibit greater bias towards the target (-) strand and be enriched in the 3’ direction relative to the PS than the first acquired spacer (+1). To test this, we independently mapped the +1 and +2 spacers in each amplicon containing at least two novel spacers (n = 1175) to characterize their respective acquisition profiles (**Figure S11**). The +1 spacer distribution closely mirrored the overall distribution shown in **Figure 3C**, while the +2 spacers showed a notable positional shift, a pattern similarly reported for priming in the type I-F CRISPR-Cas of *Pectobacterium atrosepticum* [39]. In agreement with our hypothesis, a greater proportion of +2 spacers mapped to the target (-) strand (increasing from 23.83% to 37.11%), and more were located 3’ of the PS (increasing from 12.51% to 22.64%). These positional shifts demonstrate that pre-spacer substrates are sampled sequentially and directionally during interference-driven priming, with their order of acquisition directly recorded within CRISPR arrays.

### Isolated and orphaned arrays in *Pseudomonas aeruginosa* are horizontally acquired

Isolated and orphaned CRISPR arrays can emerge when CRISPR-Cas loci associate with MGEs and are translocated within and across genomes [24]. To investigate the origin of the isolated CR3 array in PA10145 and assess its broader prevalence, we surveyed publicly available *Psa* genomes for CRISPR arrays located at loci distant from their cognate *cas* genes. From 1,198 complete whole-genome sequences, we identified 1,657 CRISPR arrays, with type I-F being the most abundant subtype (78%), followed by types I-E (18%), I-C (2%) and IV-A1 (0.5%) (**Figure 4**). Arrays located distal to their cognate *cas* genes were identified in both types I-F (n = 429) and I-E (n = 18). Of these, isolated arrays were only observed in type I-F (n = 281), with most residing at least 1 Mbp from their cognate *cas* (**Table S4**), while orphaned arrays were found in both types I-F (n = 148) and I-E (n = 18). Type I-E orphaned arrays are embedded within putative prophages, implicating MGE-mediated mobility as the mechanism by which these arrays disseminate across *Psa* genomes (**Figure S12**).

**Figure 4.**
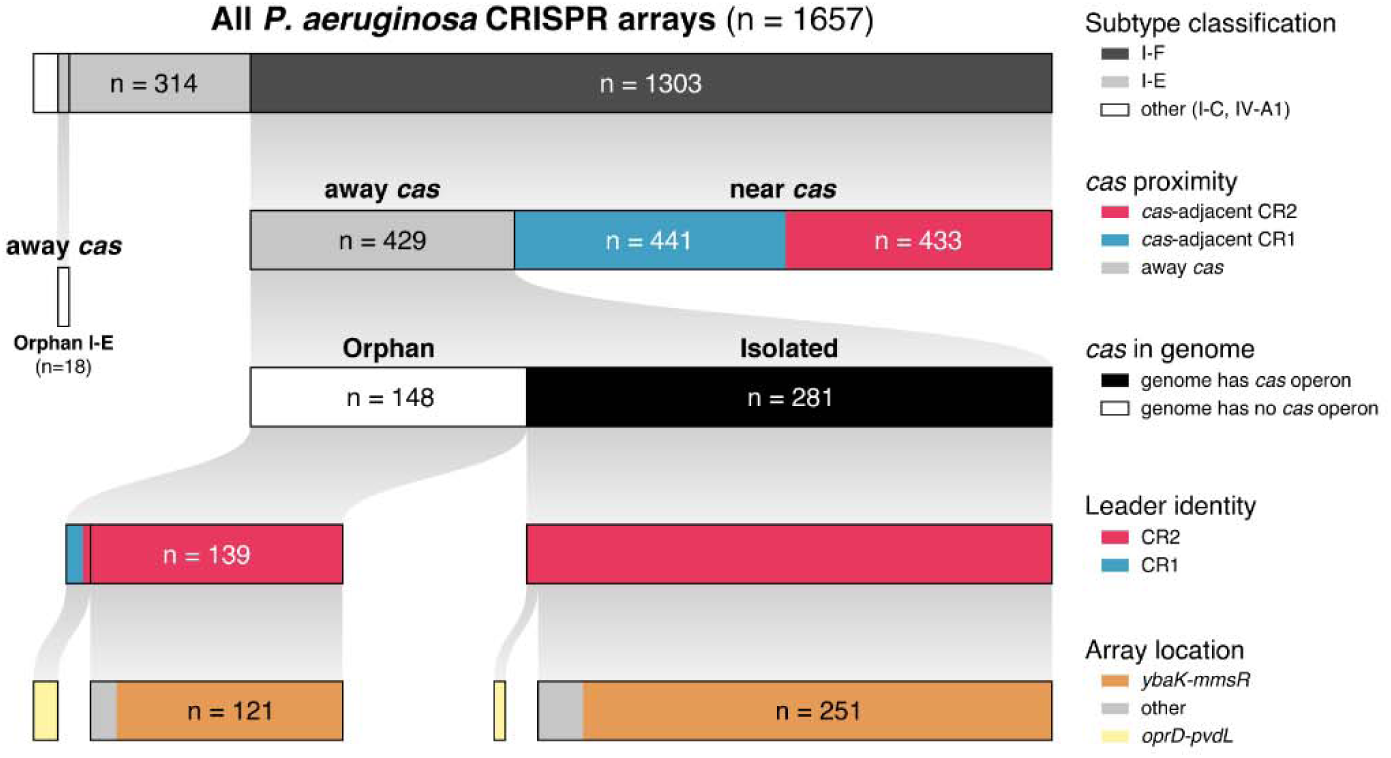
Classifications of all *Psa* CRISPR arrays. A total of 1,657 CRISPR arrays were identified from all complete NCBI-published *Psa* genomes. Type I-F was most frequent followed by I-E, I-C, then IV-A1. Only arrays of types I-F (n = 429) and I-E (n = 18) did not have an adjacent intact *cas* operon. Orphan CRISPR arrays were observed for both I-F and I-E subtypes. Isolated CRISPR arrays only occurred as type I-F; all of which have leaders that are >70% identical to the *cas*-adjacent CR2. The majority of I-F CRISPR arrays away from *cas* were located within the *ybaK-mmsR* region which is the same position as the isolated CR3 of PA10145.

All isolated and orphaned type I-F CRISPR arrays carried leader sequences that are at least 70% identical to the adjacent CR1 or CR2 arrays (**Figure 4**), with the majority bearing great similarity to CR2. Approximately ∼86% of isolated CRs and orphaned arrays occupy the same genomic neighborhood, positioned between *ybaK* and *mmsR* (**Figure 1**). Additional isolated and orphaned arrays were found outside of the *ybaK-mmsR* locus and retained CR2-like leader sequences (**Figure S13**). The remaining orphaned arrays, including all those with leader sequences more related to CR1, were located between *oprD* and *pvdL* (**Figure S14**). This is the canonical position of the type I-F CRISPR-Cas locus but instead lacked an intact *cas* operon entirely. These orphaned arrays likely represent inactive remnants of CRISPR-Cas reductive evolution [41]. The striking conservation of CR2-like leader sequences and genomic positioning across isolated and orphaned arrays throughout *Psa* genomes suggests that these elements originated through recombination-mediated translocation of *cas-*adjacent arrays to a defined loci elsewhere in the genome.

Given that the majority of isolated and orphaned CRISPR arrays occupy conserved genomic locations, their consistent positioning makes them well-suited for comparative genomic analyses aimed at resolving their evolutionary origins. In particular, the isolated CR3 may have emerged through CR2 duplication or acquired laterally via HGT. To investigate the possible role of MGEs in shaping this distribution, we analyzed the genomic neighborhoods flanking CR2, CR1, and CR3 (**Figure 5**). If MGEs have integrated at these loci, their presence would result in an expanded distance between two conserved genes defining each CRISPR neighborhood: (1) *oprD-cas1f* for CR2, (2) *cas6f-pvdL* for CR1, (3) *ybaK-mmsR* for CR3 (**Figure 1**). While most *Psa* genomes showed comparable lengths across all three regions, the *oprD-cas1f* and the *ybaK-mmsR* regions flanking CR2 and CR3, respectively, exhibited notable variability, with some genomes spanning 5- to 40-fold greater distances than typical. Crucially, all extended neighborhoods were associated with MGEs. Insertional sequences (IS) were detected near or within all CR arrays (**Figure S15**), while prophages and integrative conjugative islands (ICEs) were specifically and frequently associated with CR2 and CR3, contributing to the expanded lengths of these regions (**Figure 5**; **Figure S16-S17**). In some instances, these MGEs appear to have integrated directly within CRISPR arrays or within intergenic regions of the CRISPR-Cas operon (**Figure S15-S17**). The selective association of highly transmissible MGEs with CR2 and CR3, but not CR1, strongly supports the hypothesis that CR3 emerged through MGE-mediated translocation. Conversely, the absence of these MGEs associating with CR1 likely accounts for the lack of orphaned or isolated CR1 across *Psa* genomes.

**Figure 5.**
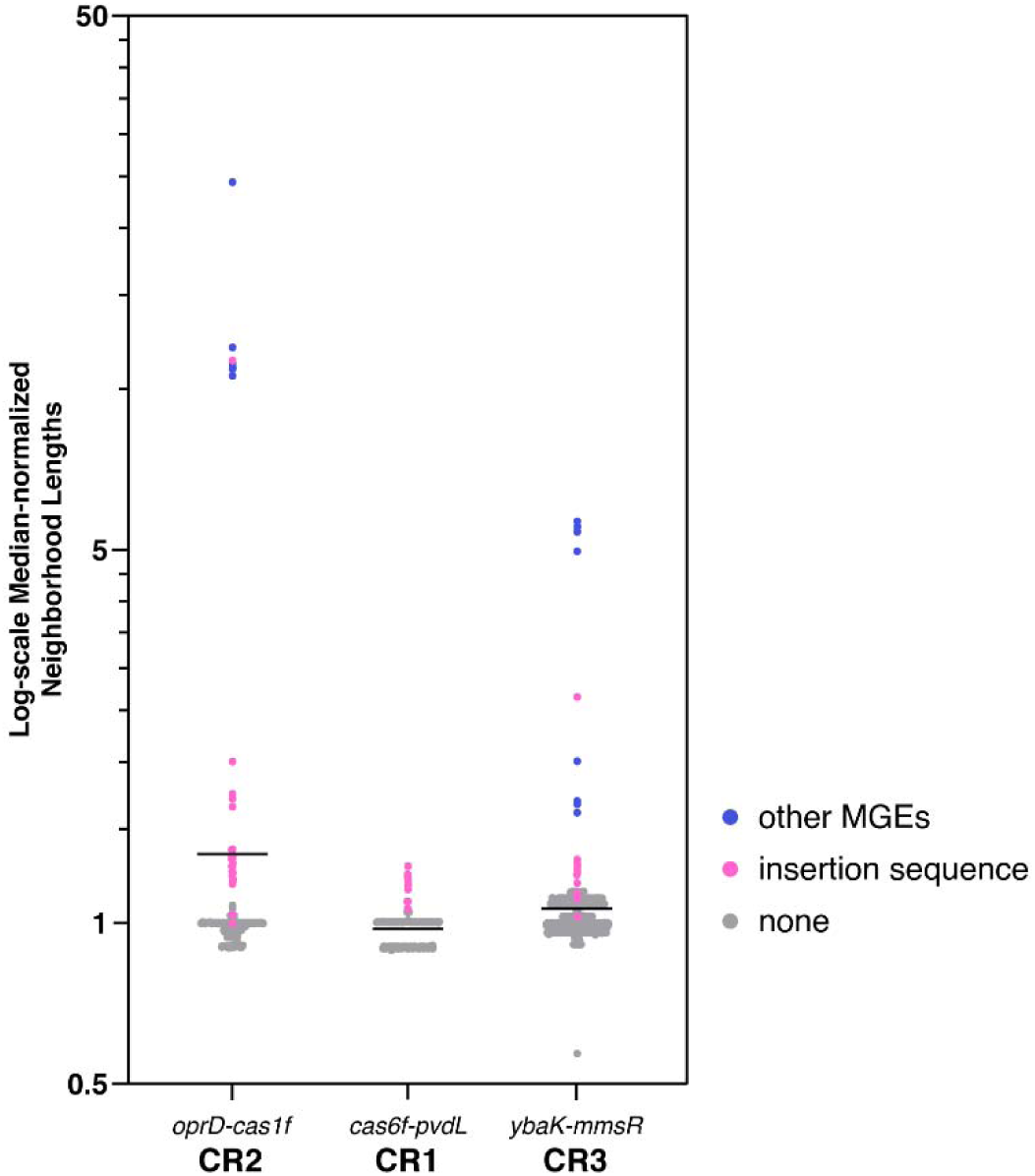
Neighborhood lengths of type I-F CRISPR arrays. Two conserved genes flanking each CRISPR array of PA10145 were used to query neighborhood expansion due to MGE insertion across *Psa* genomes. Assessed regions were *oprD-cas1f* for CR2 (n = 428), *cas6f-pvdL* for CR1 (n = 422), and *ybaK-mmsR* for CR3 (n = 1,174). CRISPR lengths were subtracted from distances prior to normalization with the median of each neighborhood length. Insertion sequences were detected among all regions. However, only the CR2 and CR3 loci contain highly transmissible MGEs such as prophages and integrative conjugative elements (ICE), resulting in considerable neighborhood expansion.

CRISPR arrays function as the genetic record of past infections, with each spacer encoding the identity of a previously encountered invader. Thus, the relatedness of spacer contents between arrays can provide insights to their likely evolutionary origins since more distal spacers represent more ancient immune encounters, whereas proximal spacers reflect more recent acquisitions. Should the isolated CR3 have emerged from a duplication event, spacers would be expected to be shared between CR2 and CR3; specifically, spacers proximal within CR2 arrays should be found at the distal end of the isolated CR3 arrays, serving as historical evidence for the duplication of a minimal CR2 leader along with its neighboring spacers. To investigate this, spacers from each CRISPR array were compiled and subjected to network analysis using CCTK [42] to visualize inter-array spacer sharing (**Figure S18**). The CR2 array network was largely interconnected yet expansive, consistent with their vertical transmission and hyperactive spacer acquisition [33,37]. CR1 arrays likewise formed an interconnected network, though comparatively more compact, reflecting its lower frequency of acquisition relative to CR2 [33,37]. In contrast, CR3 arrays exhibited largely unique spacer profiles, sharing at most a single spacer with CR2 arrays, with no overlaps found between distal CR3 spacers and proximal CR2 spacers. This limited sharing argues against a simple duplication event as the origin of the isolated CR3 and instead points to a more complex evolutionary history.

The phylogeny of CRISPR arrays can be inferred from their spacer contents and profiles through maximum parsimony [42]. A CRISPR spacer profile encompasses both the sequence identity of each spacer and the chronological order of their acquisition, reflecting the evolutionary history of the array as spacers are gained or lost over time. By comparing the phylogenies of the adjacent CR2 and CR1 arrays with that of the isolated CR3, the evolutionary trajectory of CR3 can be traced relative to its native counterparts. Since CR2 and CR1 are assumed to be vertically co-transmitted, their spacer profiles were concatenated to generate a phylogenetic tree of the type I-F *Psa* CRISPR-Cas systems based on infection history (**Figure 6**). A separate phylogenetic tree for the isolated CR3 arrays was constructed from the two largest CR3 networks, yielding five major clades corresponding to the 5 major clusters in the CR3 network (**Figure S19**). These CR3 lineages were subsequently mapped onto the base tree constructed from the infection histories of adjacent arrays to assess their evolutionary relationships (**Figure 6**). Several CR3 arrays clustered with specific clades of the base tree, representing strains which originated from a common ancestor and diverged through CRISPR adaptation. However, the overall distribution of isolated CR3 arrays across the base tree was sporadic, suggesting that these arrays have been independently acquired across diverse type I-F *Psa* CRISPR-Cas lineages which is consistent with the modular and mobile nature of isolated CRISPR arrays.

**Figure 6.**
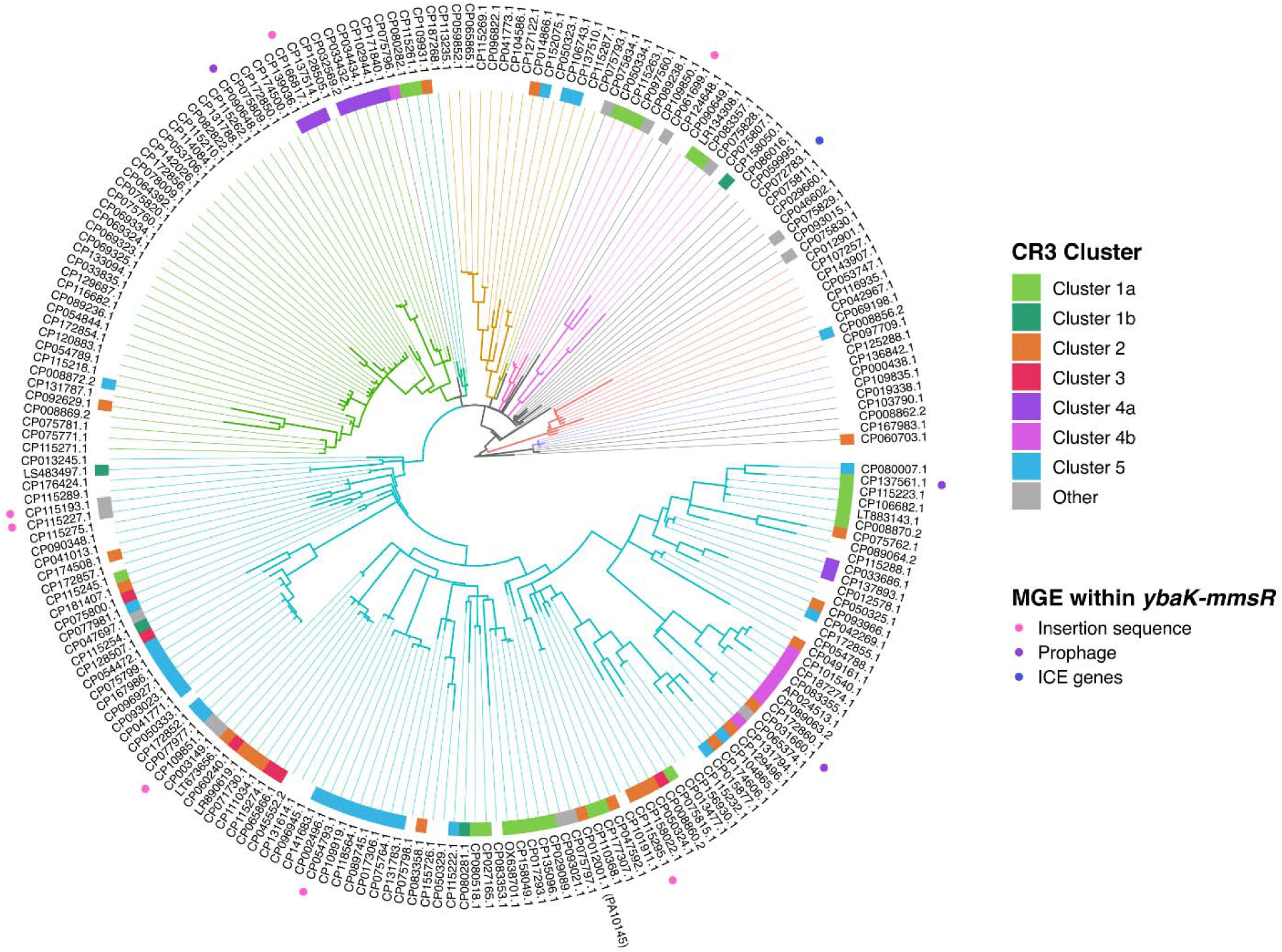
Isolated CR3 arrays are sporadically distributed across *Psa*. A maximum parsimony tree of type I-F CRISPR-Cas in *Psa* was constructed by using concatenated spacer information of the two native arrays: CR2 and CR1. Genomes with any disrupted array were excluded from tree generation. Clades grouped through hierarchical clustering have their branches colored accordingly. Plotted along the tree are the CR3 lineages of each genome as determined from the CR3 spacer sharing network and alignments. MGE insertions within the *ybaK-mmsR* locus are also indicated on each representative genome.

To further resolve the evolutionary origins of the isolated CR3 array in PA10145, genomes harboring closely related CR3 arrays were scrutinized in greater detail (**Figure 7**). Two such genomes (CP093966.1 and CP111034.1) were found to harbor nearly identical isolated CR3 arrays despite possessing phylogenetically distant adjacent arrays, indicating that these genomes share a much recent CR3 donor with respect to the evolution of their native CRISPR-Cas loci. Conversely, two genomes with highly similar infection histories in their adjacent CR2 and CR1 arrays (CP115274.1 and CP111034.1) contained isolated CR3 arrays that were phylogenetically distant from one another, further decoupling the evolutionary trajectories of isolated and adjacent arrays. These phylogenetic findings, in conjunction with the frequent association with MGEs within CR3 genomic neighborhoods, demonstrate that isolated CR3 arrays are horizontally disseminated across *Psa* genomes, likely facilitated by MGE-mediated mobilization.

**Figure 7.**
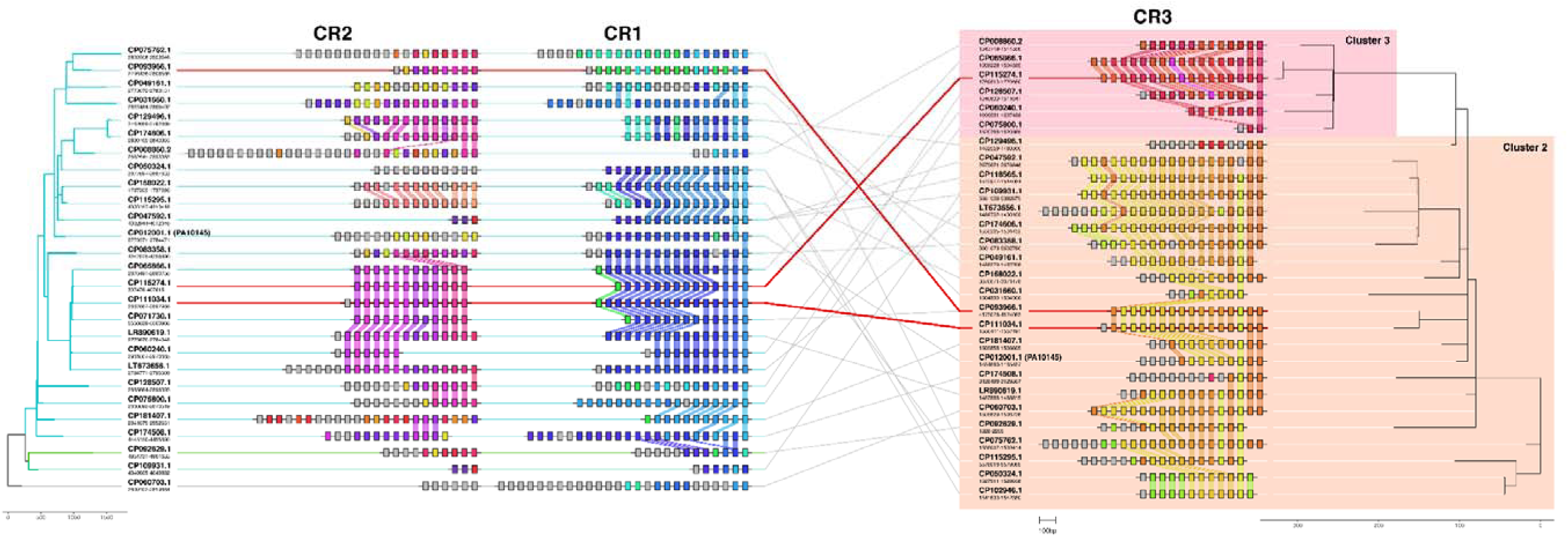
The isolated CR3 array is horizontally acquired. CR2, CR1, and CR3 of *Psa* genomes with closely related isolated arrays to PA10145-CR3 are aligned and plotted according to their respective trees. Links between native CRISPR arrays and their isolated CR3 represent 30 genomes with unique combinations of these arrays. As highlighted in red, two nearly identical isolated arrays from Cluster 3 can be harbored by two different genomes (CP093966.1 and CP111034.1) with evolutionarily distant native arrays. Conversely, two *Psa* genomes with closely related native arrays (CP115274.1 and CP111034.1) can contain entirely different CR3 arrays.

## Discussion

Isolated and orphaned CRISPR arrays are widely distributed across bacterial and archaeal genomes. These arrays are generally interpreted as vestiges of *cas* reductive evolution [24], or remnants of once-complete CRISPR-Cas systems whose effector machinery was selectively lost to permit greater horizontal gene transfer [41] or maintained as orphan arrays carrying anti-*cas* spacers that actively prevent the reacquisition of functional CRISPR-Cas systems [27]. Despite their prevalence, the function and evolutionary origin of these isolated and orphaned arrays in *P. aeruginosa* remained poorly understood. Here, we demonstrate that isolated CR3 array participates in the type I-F CRISPR-Cas immunity of PA10145 as a functional CRISPR array, reducing the transformation efficiency and plasmid maintenance of an engineered vector derived from a CR3 spacer and incorporating new spacers through interference-mediated primed adaptation. Rather than representing a vestigial remnant, the isolated CR3 array appears to be a product of duplication of the adjacent CR2 array to a distant genomic location, with the duplication encompassing the upstream region shared between CR2 and CR3, accounting for their nearly identical leader sequences. The minimal spacer shared between the isolated CR3 and adjacent CR2 arrays could be reconciled by two possible scenarios. First, the duplication event may have initially copied only a minimal CRISPR array consisting of a leader and a single repeat, which is sufficient for spacer acquisition [43]. Alternatively, isolated arrays may have lost the original co-duplicated spacers over the course of CRISPR evolution. Regardless, phylogenetic evidence from this study demonstrates that isolated arrays have evolved as separate genetic entities following their horizontal transfer to distant genomic locations, diverging from their adjacent arrays through independent spacer acquisition.

Type I-F CRISPR-Cas systems in *P. aeruginosa* typically possess two CRISPR arrays, CR2 and CR1, with CR2 exhibiting higher acquisition activities as demonstrated in the PA14 strain [33,37]. In PA10145, however, high-throughput sequencing of CRISPR arrays following primed adaptation revealed that the number of newly acquired spacers in CR2 was notably lower than in CR1. This suggested reduced acquisition activity at CR2 in PA10145, in contrast to CR2 in PA14 despite the two sharing the same leader sequence. Interestingly, the number of new spacers acquired by the isolated CR3 following primed adaptation was comparable to that of CR2, which may be attributable to their nearly identical leader sequences (**Table 1**). The co-existence of two CRISPR arrays with almost identical leaders within a genome raises the possibility that the isolated CR3 functions as an additional genetic memory bank for spacer integration, redistributing acquisition events that would otherwise be directed solely to CR2. This redistribution is further reflected in the spacer counts of both wild-type strains: CR2 and CR1 in PA14 contain 21 and 14 spacers, respectively, whereas in PA10145, these counts are reduced to 15 and 17, respectively. Broadening this analysis to all type I-F CRISPR arrays in *Psa* revealed similar acquisition dynamics among arrays with highly similar leaders co-existing within the same genome. Specifically, there was a strong negative correlation between the acquisition activities of adjacent CR2 arrays and the spacer redistribution index, defined as the degree to which an isolated array diverts spacer integration from the adjacent CR2 (**Figure S20**).

Naïve adaptation and PAM-mutant priming are inherently inefficient modes of spacer acquisition in *P. aeruginosa* [32,34]. Therefore, type I-F CRISPR-Cas systems are functionally limited when challenged with novel invaders or PAM-escape mutants. In this context, mobilization of CRISPR arrays to *cas*-harboring recipients represents a viable immunization strategy that bypasses the requirement of prior exposure to a given invader. Upon transfer, interference-proficient spacers within the acquired array can immediately confer immunity and drive rapid strand-biased primed adaptation, permanently updating the native CRISPR arrays of the recipient cell and reinforcing individual immunity. The near-identical leader sequences of CR2 and the isolated CR3 further ensure that newly acquired spacers are distributed equally across both arrays, so that the isolated array is updated simultaneously during primed adaptation. Upon subsequent horizontal transfer to naïve cells, the isolated array serves as a conduit of expanded genetic memory: providing recipient cells with a continuously evolving spacer repertoire, enabling primed adaptation to further update their native arrays, and administering the dissemination of immunological genetic memory across the bacterial population. This continuous cycle of acquisition, updating, and horizontal dissemination positions isolated CRISPR arrays as key mediators of population-level pan-immunity, enabling bacterial populations to collectively and dynamically respond to changing infectious threats through a shared pool of mobilizable genetic memory without sacrificing the genomic plasticity afforded by ongoing HGT.

The sporadic yet recurrent presence of isolated CRISPR arrays as portable genetic modules across *P. aeruginosa* genomes aligns with the “pan-immune system” model [44]. This model reconciles two seemingly contradictory observations in prokaryotic immunity: that individual bacteria frequently encode multiple defense systems acquired through horizontal gene transfer, suggesting clear fitness benefits; yet these same systems are just as frequently lost on short evolutionary timescales, implying an associated fitness burden. No single bacterium can bear the cumulative fitness cost of maintaining all possible defense systems simultaneously; instead, HGT serves as a dynamic mechanism through which individual cells can transiently access the broader defensive repertoire distributed across closely related strains [44].

The exclusivity of isolated arrays to the type I-F subtype in *P. aeruginosa* is unlikely to be coincidental. As the most prevalent CRISPR subtype in this species, isolated type I-F arrays are disproportionately likely to encounter *cas*-harboring recipient cells upon mobilization, maximizing the probability that a transferred array will be functional in a new genomic context. Within this framework, isolated arrays represent a particularly efficient class of transferable immune modules, conferring interference-competent genetic memory upon recipient cells without the fitness burden of acquiring an entire CRISPR-Cas system. Furthermore, with growing evidence that complete immune systems, including entire CRISPR-Cas loci, can themselves be horizontally transferred [11,45–47], the heterogeneous distribution of *P. aeruginosa* genomes that lack or possess type I-F CRISPR-Cas systems and isolated arrays may reflect a population-level balancing of evolutionary trade-offs [12]. Strains maintaining robust CRISPR-Cas immunity benefit from broad phage resistance but incur the cost of restricting horizontal gene transfer and limiting their capacity for rapid genomic diversification. Conversely, strains that have lost their CRISPR-Cas systems sacrifice targeted immunity but gain unrestricted access to HGT as a driver of genomic evolution and adaptation. The coexistence of these genomic states within the *P. aeruginosa* population may therefore represent a bet-hedging strategy, wherein the pangenome collectively sustains both immunological robustness and evolutionary flexibility with isolated CRISPR arrays serving as portable bridges between these two states, capable of rapidly immunizing naïve recipients while remaining compatible with ongoing genomic exchange.

The mobilization of isolated arrays for CRISPR-mediated pan-immunity may not be exclusive to *P. aeruginosa* as other bacterial species have been demonstrated to harbor CRISPR arrays distantly located from their cognate *cas*. *Yersinia pestis* has been documented to possess a third CRISPR array (YP3) also located 1 Mbp away from its cognate *cas* genes [25]. While its leader sequence significantly differs from *cas*-adjacent arrays, spacer profiles within YP3 can vary across strains which suggests its participation in CRISPR adaptation by independently acquiring spacers [25]. In an extensive investigation on all CRISPR arrays in eubacteria, multiple distal type I-F arrays with highly similar leader sequences were identified in *Xanthomonas albilineans* genomes. Type I-F arrays of *X. albilineans* can possess self-targeting spacers yet, paradoxically, its type I-F CRISPR-Cas system was actively expressed and remained functional [48]. Lethal autoimmunity was prevented by anti-CRISPR (Acr) proteins encoded by prophage regions. Although deleterious self-targeting is prevented by these Acrs, DNA binding to a chromosomal target by type I-F Cascade was still permitted, incurring gene repression of the target and implicating a potential regulatory role of isolated CRISPR arrays in gene expression [48].

## Methods

### Strains and growth conditions

*E. coli* DH5α and *Psa* ATCC 10145 used in this study were incubated at 37°C in Luria-Bertani (LB) broth with shaking at 225 rpm. Cultures on solid media are displayed on 1.5% w/v LB agar (LBA) plates. When required, 100 µg ml–1 ampicillin, 50 µg ml–1 streptomycin (1X), and/or 1 mM isopropyl-β-D-thiogalactopyranoside (IPTG) was supplemented in the media.

### Plasmid construction

*cas*, CR2, CR1, and CR3 UIRs were amplified from a PA10145 colony then directionally cloned to a promoterless *lacZ* in pSEVA445 to create transcriptional *lacZ* gene fusion constructs reporting the promoter activities of CRISPR-Cas. mCherry inserts were PCR amplified from pQE-80L-mCherry, digested using BamHI and SalI, then ligated to linearized pTA100. The criteria for choosing proximal spacers to be incorporated within the reverse mCherry primers were based on sequence complexity. The second spacer of CR2 and the third spacers of CR1 and CR3 were appended. T plasmid inserts contain a protospacer from each CRISPR array and a canonical 3’-GG PAM while NT plasmid inserts possess a mutated 3’-GT PAM. Plasmid extracts from pink colonies were then sent for DNA sequencing with primers that flank the MCS of pTA100 (SML084+SML085). However, pTA100 was found to be incompatible with PA10145. Thus, mCherry inserts were subcloned to pSEVA451 which contains a broad-host range *ori*: RSF1010. All plasmids with inserts constructed in this study have been sequence verified.

### Heat shock transformation of PA10145

Chemical competency in *Psa* was induced as previously described [49]. One microgram of each engineered construct alongside the control no PS vector was added and mixed with the competent cells by pipetting. Cells were then heat shocked and recovered prior to display on 15X Sm LBA. Colonies were counted using ImageJ then transformation efficiency was calculated.

### 4-Methylumbelliferyl-β-D-galactopyranoside (MUG) assay

Overnight cultures of PA10145 containing pSEVA445 constructs with *lacZ* fused to type I-F CRISPR-Cas UIRs were prepared in biological duplicates along with an empty pSEVA445 as control. These were then subcultured from a starting OD_600_ of 0.05 then grown for an hour. Three aliquots per culture were obtained as technical triplicates to the 96-well master plate, freeze-thawed, then transferred to the assay plate. MUG reagent was added per well then measured using EnSight Multimode Plate Reader (PerkinElmer) with excitation and emission wavelengths set at 365 nm and 455 nm, respectively. Readouts were normalized to the measured OD_600_.

### Plasmid maintenance assay

At day 0, 24-h cultures consisting of a single colony of PA10145, transformed with T or NT plasmids, were generated in triplicate with selection at 15X Sm alongside PA10145 with no PS plasmid as control. Day 0 inoculum was washed prior to subculture without selection as the day 1 culture. Subsequent 24-h cultures were passaged from the previous culture until day 5. Cultures were then pelleted to obtain a sample for assessing CRISPR expansion at the populational level.

In parallel, dilutions were created until 10^-5^ then displayed on plates with selection at day 0. For subsequent subcultures, plates without selection were generated from the 10^-6^ dilution. Dilution plates were incubated for 12-16 hours at 37°C. One hundred (100) isolated colonies from each plate were replica-plated onto non-selective (NS) and selective (S) LBA. The number of surviving colonies on the S plate corresponded to percent plasmid maintenance. Three random antibiotic-sensitive colonies per NS plate were assessed for CRISPR expansion.

### Detection of CRISPR expansion

PCR templates are obtained from the pooled populations of day 5 cultures or from three random sensitive colonies from each NS plate at day 5. Primer pairs bind to the UIR and the third spacer of CR2 (SML507-SML508), CR1 (SML510-511), and CR3 (SML507-SML509). Reactions were run in AGE at 100V for 35 minutes in 2% agarose gel.

### High-throughput spacer acquisition analysis

CRISPR arrays were PCR-amplified from populations 5 days post-exposure to the T plasmid. Bands higher than 200 bp were gel excised, extracted, then pooled according to the PS of the T plasmid set-up prior to purification. DNA were then mixed in equimolar amounts for library preparation as instructed by Oxford Nanopore Technologies (ONT). The library was then loaded to the SpotON flow cell in MinION Mk1B. Custom Python scripts were then executed to parse through the sequencing data by filtering reads that contain two or more repeats and mapping each spacer flanked by two repeats to a position on the backbone of the T plasmid. Each novel spacer was also binned according to which CRISPR had acquired it based on the two pre-existing spacers included within the read.

### CRISPR analysis of all *Psa* arrays

The complete genomic DNA sequence of PA10145 was obtained from the NCBI database under the GenBank Accession No. CP012001.1. The genome was then inputted to CRISPRCasTyper v1.8.0 [50] for annotation. Within PA10145, repeats were visualized using the RNAfold webserver [51]. Upstream sequences of each of the three CRISPR arrays were aligned with Clustal Omega [52] and visualized in ESPript [53].

Complete *Psa* assemblies were obtained through the ncbi-datasets v16.24.0 package (accessed May 4, 2025) then inputted to CRISPRCasTyper [50] to classify all *Psa* CRISPR arrays by subtype and determine CRISPR proximity to an intact *cas* operon. Leader identities of all I-F arrays were obtained by aligning a 200 bp upstream sequence of each array to the leaders of *cas*-adjacent CR2 and CR1 in PA10145. Given the similar relative genomic positions of I-F CRISPR arrays, gene neighborhoods of all arrays were measured according to base-pair length using *oprD-cas1f* for CR2, *cas6f-pvdL* for CR1, and *ybaK-mmsR* for CR3. Should a region contain an array, the CRISPR length was subtracted from the neighborhood distance as normalization. Regions were re-annotated on Prokka v1.14.6 [54] for uniformity in annotation and detection of insertion sequences (IS). Additional MGE identification was performed through the PHASTEST [55] and ICEberg [56] webservers for prophages and integrative conjugative elements (ICE), respectively. Representative sequences were then visualized through clinker v0.0.29 [57].

The spacer profiles of all I-F *Psa* CRISPR arrays were analyzed using the CRISPR Comparison Toolkit v1.0.3 (CCTK) [42]. Relationships of I-F CRISPR arrays within each subset (CR2, CR1, and CR3) are represented in maximum parsimony trees through CRISPRtree. To generate a tree depicting the phylogenies of native *Psa* CRISPR arrays, spacer information of *cas*-adjacent CR2 and CR1 of each genome were concatenated prior to tree construction, excluding genomes with any disrupted CRISPR. Generated networks and trees were visualized in R v4.5.1 using the igraph v2.2.0 [58] and ggtree v3.16.3 [59] packages, respectively. Hierarchical clustering for grouping related clades was performed using the dendextend v1.19.1 [60] and phylogram v2.1.0 [61] packages. Cophylogenetic trees were generated using the phytools v2.5-2 [62] package. Spacer alignments were then visualized through clinker [57].

