## Supplementary material for "Acquisition of novel arrays via horizontal gene transfer rewire CRISPR-mediated defense in *Pseudomonas aeruginosa*": Table S1 - Table S7

### Supplemental Tables

**Table S1. Representative top hits of CR2 spacers.** Hits scored by nucleotide match/mismatch (+1/-1) and I-F PAM match (+5).

| **CR2-sp#** | **Accession** | **Identity** | **Description** | **Score** | **Start** | **End** |
| --- | --- | --- | --- | --- | --- | --- |
| 1 | OQ572410 | phage | *Pseudomonas* phage Y8 | 37 | 4143 | 4174 |
| 2 | CP093026.1 | plasmid | *Pseudomonas aeruginosa* H06 plasmid unnamed2 | 37 | 2950 | 2981 |
| 3 | CP093027.1 | plasmid | *Pseudomonas aeruginosa* H06 plasmid unnamed3 | 37 | 3702 | 3733 |
| 4 | OP966821 | phage | *Pseudomonas* phage Fyn8 | 37 | 29001 | 29032 |
| 5 | NC_073672.1 | phage | *Pseudomonas* phage PA8 | 37 | 42793 | 42824 |
| 6 | OQ354711 | phage | *Pseudomonas* phage WX_Y | 37 | 6747 | 6716 |
| 7 | CP093026.1 | plasmid | *Pseudomonas aeruginosa* H06 plasmid unnamed2 | 37 | 3034 | 3003 |
| 8 | CP012579.1 | prophage | *Pseudomonas aeruginosa* PA_D5 (1513117-1520871) | 35 | 4723 | 4754 |
| 9 | CP092031.1 | plasmid | *Pseudomonas aeruginosa* ZS-PA-05 plasmid pZS-PA-05 | 37 | 16390 | 16359 |
| 10 | NC_027992.1 | phage | *Pseudomonas* phage JBD25 | 37 | 34381 | 34412 |
| 11 | WTXS01000205.1 | plasmid | *Pseudomonas aeruginosa* JX05 plasmid unnamed1 | 37 | 2009 | 2040 |
| 12 | MZ773939 | phage | *Pseudomonas* phage PP9W | 35 | 46892 | 46923 |
| 13 | OP709960 | phage | *Pseudomonas* phage zjk6 | 37 | 26208 | 26239 |
| 14 | CP012582.1 | prophage | *Pseudomonas aeruginosa* PA_D21 (1513116-1520870) | 37 | 3844 | 3813 |
| 15 | OR683411 | phage | *Pseudomonas* phage Solano | 37 | 11832 | 11863 |

**Table S2. Representative top hits of CR1 spacers.** Hits scored by nucleotide match/mismatch (+1/-1) and I-F PAM match (+5).

| **CR1-sp#** | **Accession** | **Identity** | **Description** | **Score** | **Start** | **End** |
| --- | --- | --- | --- | --- | --- | --- |
| 1 | CP015001.1 | island | *Pseudomonas aeruginosa* PA1088 (3280318-3294459) | 33 | 12035 | 12004 |
| 2 | PQ758391 | phage | *Pseudomonas* phage 5a-Dazarov | 33 | 3693 | 3725 |
| 3 | WTXS01000205.1 | plasmid | *Pseudomonas aeruginosa* JX05 plasmid unnamed1 | 37 | 1740 | 1771 |
| 4 | WTXS01000001.1 | plasmid | *Pseudomonas aeruginosa* JX05 plasmid pBH6 | 37 | 117329 | 117298 |
| 5 | NO HIT |  |  |  |  |  |
| 6 | CP115212.1 | plasmid | *Pseudomonas aeruginosa* F064 plasmid pF064 | 35 | 7456 | 7487 |
| 7 | IMGVR | viral | UViG_2700989440_000001 (2700989440-2701063904) | 35 | 1828 | 1859 |
| 8 | CP124661.1 | plasmid | *Pseudomonas aeruginosa* 2022CK-00160 plasmid unnamed1 | 37 | 2226 | 2195 |
| 9 | NO HIT |  |  |  |  |  |
| 10 | OP709960 | phage | *Pseudomonas* phage zjk6 | 35 | 21467 | 21436 |
| 11 | NC_030931.1 | phage | *Pseudomonas* phage phi2 | 37 | 16387 | 16418 |
| 12 | PP869205 | phage | *Pseudomonas* phage PA_L9 | 35 | 84871 | 84902 |
| 13 | NC_030931.1 | phage | *Pseudomonas* phage phi2 | 38 | 16027 | 16059 |
| 14 | OQ594956 | phage | *Pseudomonas* phage vB_Pae_LESphi3 | 37 | 4600 | 4569 |
| 15 | OK041467 | phage | *Pseudomonas* phage PAE2 | 37 | 18481 | 18450 |
| 16 | NC_017549.1 | prophage | *Pseudomonas aeruginosa* NCGM2.S1 (3075868-3096936) | 35 | 5241 | 5210 |
| 17 | NO HIT |  |  |  |  |  |

**Table S3. Representative top hits of CR3 spacers.** Hits scored by nucleotide match/mismatch (+1/-1) and I-F PAM match (+5).

| **CR3-sp#** | **Accession** | **Identity** | **Description** | **Score** | **Start** | **End** |
| --- | --- | --- | --- | --- | --- | --- |
| 1 | KM389229 | phage | *Pseudomonas* phage F_TK1718spPAK | 37 | 19922 | 19953 |
| 2 | KM389229 | phage | *Pseudomonas* phage F_TK1718spPAK | 37 | 18856 | 18825 |
| 3 | CP014948.1 | prophage | *Pseudomonas aeruginosa* N17-1 (3096395-3107918) | 37 | 1755 | 1724 |
| 4 | CP093025.1 | plasmid | *Pseudomonas aeruginosa* H06 plasmid unnamed1 | 37 | 2861 | 2892 |
| 5 | OP709960 | phage | *Pseudomonas* phage zjk6 | 32 | 23949 | 23918 |
| 6 | CP092847.1 | plasmid | *Pseudomonas aeruginosa* ISS SRV-K plasmid pSRV_K_1 | 33 | 26874 | 26843 |
| 7 | MK034952 | phage | *Pseudomonas* phage Dobby | 37 | 32419 | 32388 |
| 8 | CP020602.1 | plasmid | *Pseudomonas aeruginosa* E6130952 plasmid pJHX613 | 37 | 31255 | 31224 |
| 9 | NO HIT |  |  |  |  |  |
| 10 | NC_008357.1 | plasmid | *Pseudomonas aeruginosa* E6130952 plasmid pJHX613 | 35 | 33310 | 33279 |
| 11 | CP137006.1 | plasmid | *Citrobacter koseri* K1219 plasmid pK1219-2 | 35 | 18971 | 19002 |
| 12 | KM389412 | phage | *Pseudomonas* phage F_HA1961sp | 37 | 70 | 39 |
| 13 | PQ287282 | phage | *Pseudomonas* phage Komp_PA14_gP | 37 | 42450 | 42419 |

**Table S4. Distances of I-F isolated arrays from their cognate *cas* operons.** Genome circularization was taken into account.

| **Distance from *cas* (bp)** | **Count** | **Percentage** |
| --- | --- | --- |
| 30,000 – 500,000 | 15 | 5.34% |
| 1,000,000 – 1,500,000 | 225 | 80.07% |
| 1,500,000 – 2,000,000 | 35 | 12.46% |
| 2,000,000 – 2,500,000 | 3 | 1.07% |
| 2,500,000 – 3,000,000 | 1 | 0.36% |
| 3,000,000 – 3,500,000 | 2 | 0.71% |

**Table S5. Strains used in this study**

| **Strain** | **Genotype/Phenotype** | **Reference** |
| --- | --- | --- |
| *Escherichia coli* DH5α | F-, Φ80*lac*ZΔM15Δ(*lac*ZYAargF) U169 *rec*A1 *end*A1 *hsd*R17 (rK-, mK+) *pho*A *sup*E44 λ– *thi*-1 *gyr*A96 *rel*A1 | Gibco/BRL |
| *Pseudomonas aeruginosa* ATCC 10145 | WT | ATCC |

**Table S6. Plasmids used in this study**

| **Plasmid** | **Description** | **Primers** | **Plasmid Backbone** | **Reference** |
| --- | --- | --- | --- | --- |
| pQE-80L-  mCherry | IPTG-inducible mCherry expression vector, high copy number, ColE1 *ori*, Amp^R^ |  |  | (Richter et al., 2014) |
| pTA100 | pQE-80L derivative with Sm/Sp resistant cassette, Sm/Sp^R^, ColE1 *ori* |  |  | (Fineran et al., 2009) |
| pSEVA451 | MCS-default cargo, Sm/Sp^R^, RSF1010 *ori* |  |  | (Martínez-García et al., 2020) |
| pSML0067 | Vector harboring no PAM or protospacer with mCherry reporter | SML465+SML466 | pTA100 | This study |
| pSML0068 | Vector harboring PAM+CR2_sp2 with mCherry reporter | SML465+SML497 | pTA100 | This study |
| pSML0069 | Vector harboring PAM+CR1_sp3 with mCherry reporter | SML465+SML498 | pTA100 | This study |
| pSML0070 | Vector harboring PAM+CR3_sp3 with mCherry reporter | SML465+SML499 | pTA100 | This study |
| pSML0071 | Vector harboring mutPAM+CR2_sp2 with mCherry reporter | SML465+SML500 | pTA100 | This study |
| pSML0072 | Vector harboring mutPAM+CR1_sp3 with mCherry reporter | SML465+SML501 | pTA100 | This study |
| pSML0073 | Vector harboring mutPAM+CR3_sp3 with mCherry reporter | SML465+SML502 | pTA100 | This study |
| pSML0101 | Vector harboring no PAM or protospacer | SML465+SML466 | pSEVA451 | This study |
| pSML0102 | Vector harboring PAM+CR2_sp2 | SML465+SML497 | pSEVA451 | This study |
| pSML0103 | Vector harboring PAM+CR1_sp3 | SML465+SML498 | pSEVA451 | This study |
| pSML0104 | Vector harboring PAM+CR3_sp3 | SML465+SML499 | pSEVA451 | This study |
| pSML0105 | Vector harboring mutPAM+CR2_sp2 | SML465+SML500 | pSEVA451 | This study |
| pSML0106 | Vector harboring mutPAM+CR1_sp3 | SML465+SML501 | pSEVA451 | This study |
| pSML0107 | Vector harboring mutPAM+CR3_sp3 | SML465+SML502 | pSEVA451 | This study |

**Table S7. Primers used in this study**

| **Primer** | **Sequence (5’-3’)** | **Description** | **RE** |
| --- | --- | --- | --- |
| **Cloning Primers** | | | |
| SML465 | TTT**GGATCC**GTGAGCAAGGGCGAGGAGG | F, binds to the 2nd codon of the mCherry gene in pQE80L-mCherry | BamHI |
| SML466 | TTT**GTCGAC**GGTCTCTCTACTTGTACAGCTCGTCC | R, no PAM or protospacer, binds downstream of the mCherry gene in pQE80L-mCherry | SalI |
| SML497 | TTT**GTCGAC**TGAAAAACCTGAAAAAACTGTTCGTTCGTGGTGGTTTGGTCTCTCTACTTGTACAGCTC | R, PAM+CR2_sp2, binds downstream of the mCherry gene in pQE80L-mCherry | SalI |
| SML498 | TTT**GTCGAC**GTTTTCACCGATGGTCATGGTCTCTTTCCCGAGGTTTGGTCTCTCTACTTGTACAGCTC | R, PAM+CR1_sp3, binds downstream of the mCherry gene in pQE80L-mCherry | SalI |
| SML499 | TTT**GTCGAC**TTGATCAGCAGTAAGAAAGGCACTCTTTATTTGGTTTGGTCTCTCTACTTGTACAGCTC | R, PAM+CR3_sp3, binds downstream of the mCherry gene in pQE80L-mCherry | SalI |
| SML500 | TTT**GTCGAC**TGAAAAACCTGAAAAAACTGTTCGTTCGTGGTTGTTTTGGTCTCTCTACTTGTACAGCTC | R, mutPAM+CR2_sp2, binds downstream of the mCherry gene in pQE80L-mCherry | SalI |
| SML501 | TTT**GTCGAC**GTTTTCACCGATGGTCATGGTCTCTTTCCCGATGTTTTGGTCTCTCTACTTGTACAGCTC | R, mutPAM+CR1_sp3, binds downstream of the mCherry gene in pQE80L-mCherry | SalI |
| SML502 | TTT**GTCGAC**TTGATCAGCAGTAAGAAAGGCACTCTTTATTTTGTTTTGGTCTCTCTACTTGTACAGCTC | R, mutPAM+CR3_sp3, binds downstream of the mCherry gene in pQE80L-mCherry | SalI |
| **Screening/Sequencing Primers** | | | |
| SML084 | TCGTCTTCACCTCGAGAAATC | F, Seq primer for pTA100 | - |
| SML085 | GTCATTACTGGATCTATCAACAGG | R, Seq primer for pTA100 | - |
| SML227 | CGCCAGGGTTTTCCCAGTCACGAC | F, pSEVA sequencing primer (taken from SEVA annotation), flanks the MCS | - |
| SML228 | AGCGGATAACAATTTCACACAGGA | R, pSEVA sequencing primer (taken from SEVA annotation), flanks the MCS | - |
| SML507 | CTTCAAATGGTTATAGGTTTTCGG | F, binds upstream of CR2 and CR3 arrays for screening CRISPR expansion | - |
| SML508 | CGTGAAAGATGACCGACTGTTT | R, binds CR2_sp2 for screening CRISPR expansion | - |
| SML509 | TTGATCAGCAGTAAGAAAGGCACC | R, binds CR3_sp3 for screening CRISPR expansion | - |
| SML510 | AGGTTGATGGTTTTTGGGTCTA | F, binds upstream CR1 array for screening CRISPR expansion | - |
| SML511 | GTCATGGTCTCTTTCCCGA | R, binds CR1_sp3 for screening CRISPR expansion | - |
