## Supplementary material for "Acquisition of novel arrays via horizontal gene transfer rewire CRISPR-mediated defense in *Pseudomonas aeruginosa*": Figure S1 - Figure S20

### Supplemental Figures

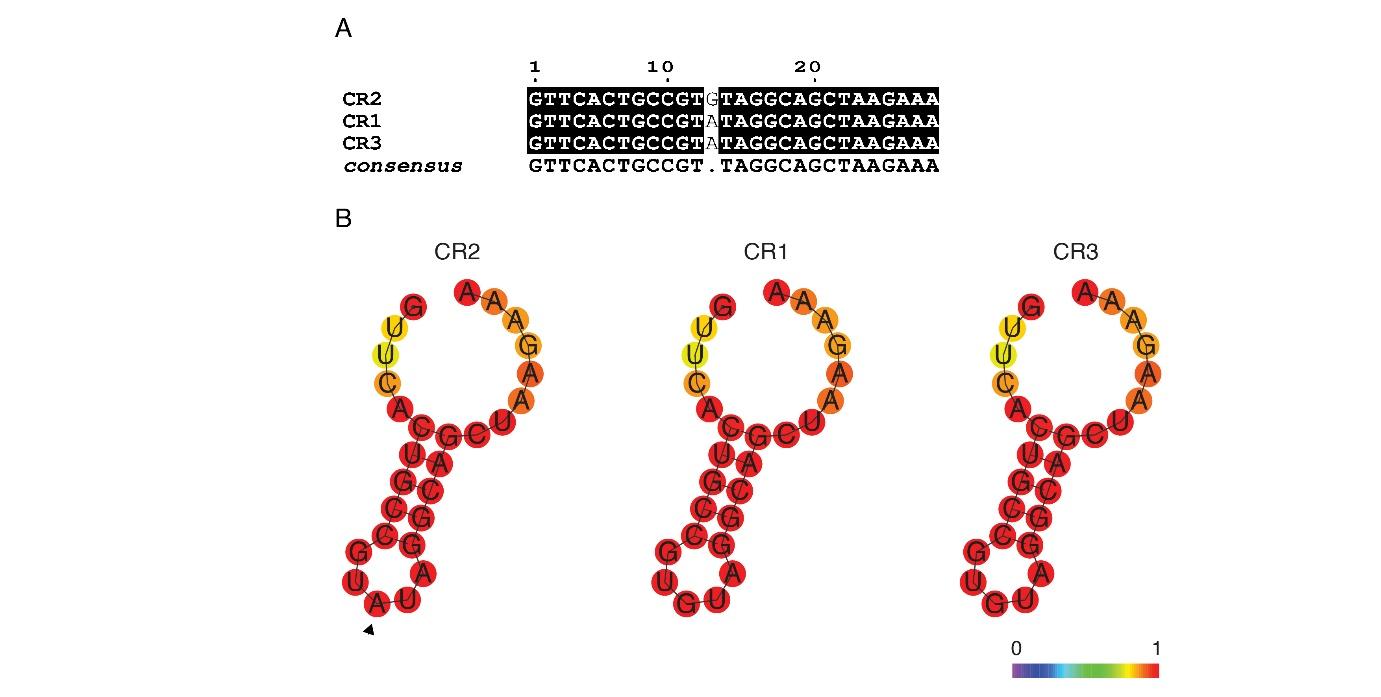

**Figure S1. Repeats of the isolated CR3 exhibit canonical RNA folding.** (A) Alignment of repeats within all three CRISPR arrays of PA10145 as visualized using ESPript [[53]](https://www.zotero.org/google-docs/?DJH9Nb). Sequences of the CR1 and CR3 repeats are identical while the CR2 repeat differs at a single base. (B) Secondary structure formation was determined by RNAfold webserver [[51]](https://www.zotero.org/google-docs/?NCGE5Z). Each of the repeats is consistent among all CRISPR arrays and matches the RNA folding found in PA14, indicating that the repeats of the isolated CR3 would be properly recognized by Cas6f leading to CR3 pre-crRNA processing into mature crRNA [[31]](https://www.zotero.org/google-docs/?Aeh8qI). CR2 contains a single base at the loop, pointed out by the black triangle. Legend at the lower right indicates the base-pair probabilities according to color. As for unpaired bases, the color represents the probability of being unpaired.

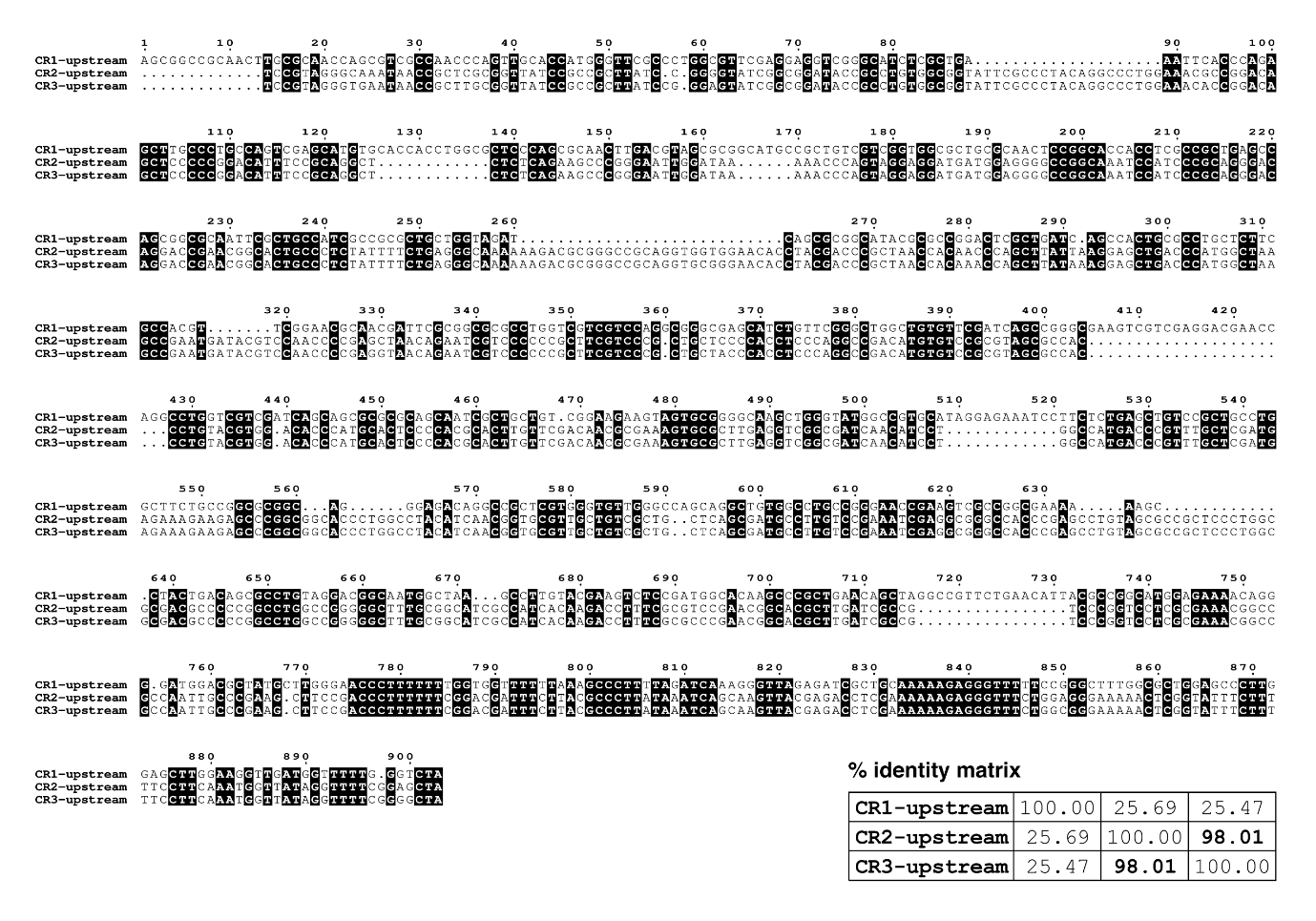

**Figure S2. Multiple sequence alignment of upstream regions of each CRISPR array of PA10145.** Clustal Omega [[52]](https://www.zotero.org/google-docs/?R4raIx) alignment as visualized in ESPript [[53]](https://www.zotero.org/google-docs/?fNa1mg). Percent identity matrix is shown on the lower right, showing a high degree of sequence similarity between the upstream ~900 bp regions between CR2 and CR3.

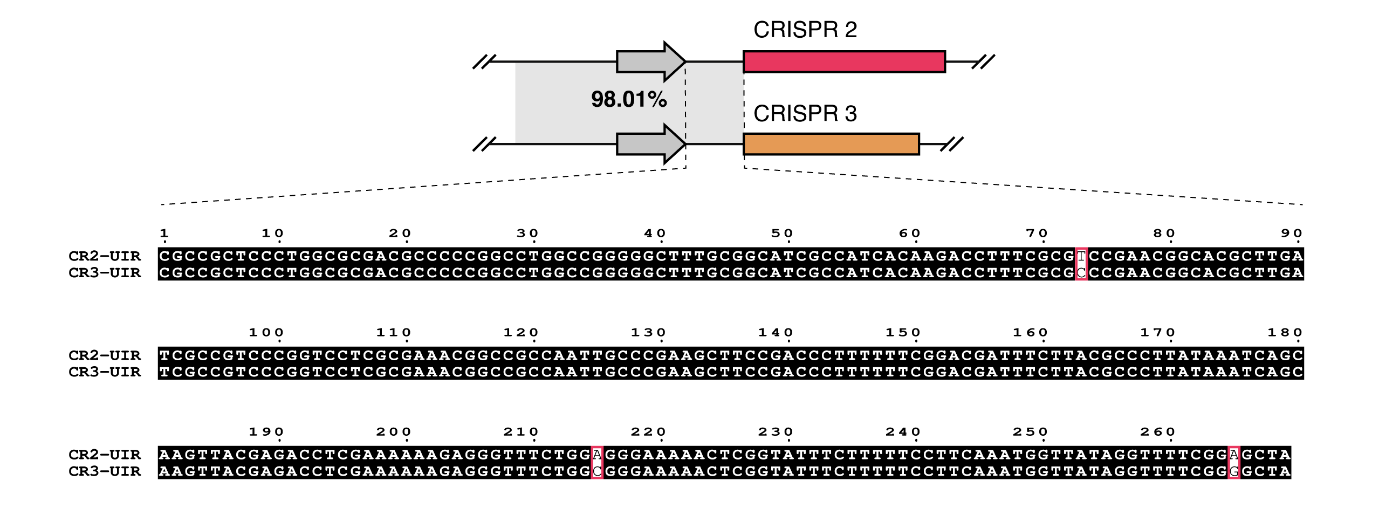

**Figure S3. The upstream intergenic regions (UIRs) of CR2 and CR3 are almost identical.** The 900 bp region upstream of CR2 and CR3 are 98.01% identical. Specifically, within the UIR, the sequences only differ in three bases as highlighted in red boxes visualized using ESPript [[53]](https://www.zotero.org/google-docs/?cCRhdd).

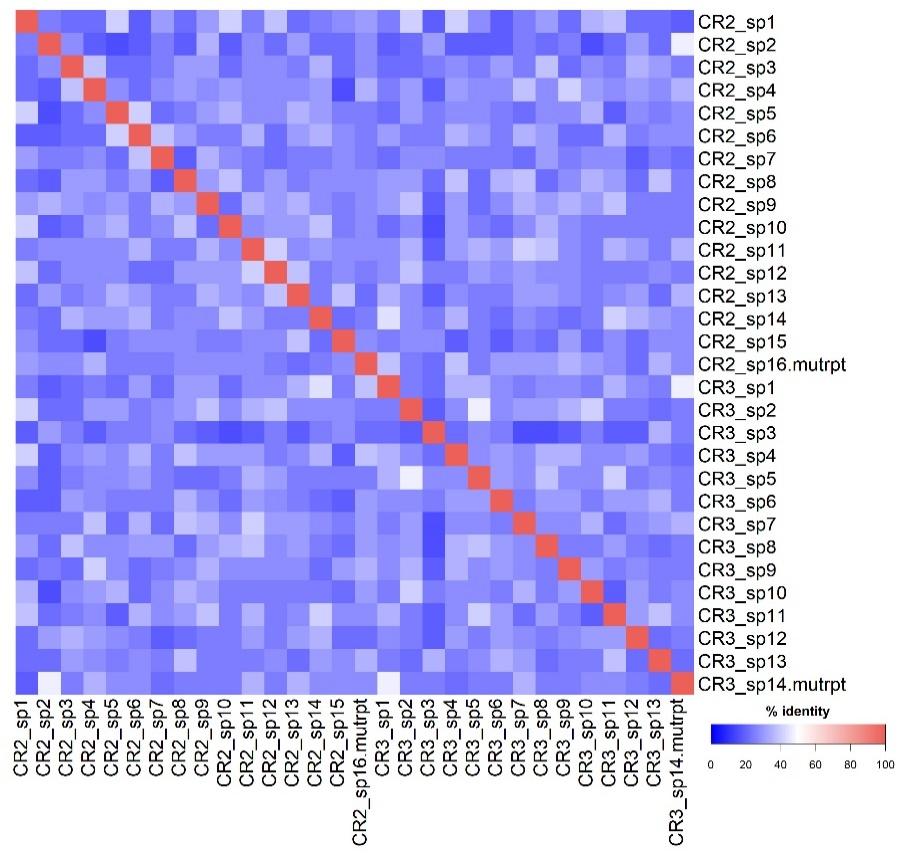

**Figure S4. The isolated CR3 has distinct spacers from CR2.** Heatmap visualization of sequence identity matrix of all spacers within CR3 and CR2 was generated by Clustal Omega [[52]](https://www.zotero.org/google-docs/?Nm32jy). The most distal spacer with a mutated repeat (mutrpt) was also included which could possibly be a remnant of incomplete excision of the repeat along with the CRISPR leader.

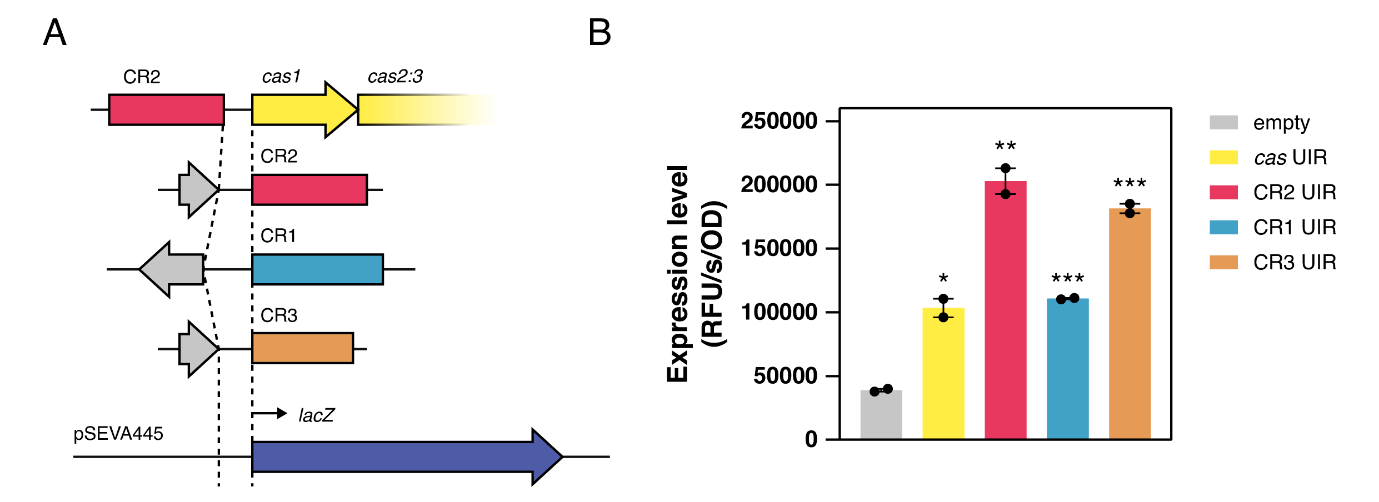

**Figure S5. The UIRs of the type I-F CRISPR-Cas system of PA10145 contain active promoters.** (A) Inserts of the upstream intergenic regions (UIRs) of each CRISPR-Cas component were PCR amplified from PA10145 then directionally cloned to the MCS of pSEVA445 which contains a promoterless *lacZ* transcriptional reporter. (B) β-galactosidase activity was measured in RFU and normalized to OD_600_ in biological duplicates. Promoter activities of the *cas*, CR2, CR1, and CR3 UIRs were significant relative to control after an hour of growth (p=0.0126, p=0.0039, p=0.0003, and p=0.0007, respectively). two-tailed Student’s t-test; ***p<0.001; **p < 0.01; *p < 0.05. Significance directly above experimental bars indicates comparison to the empty control. Bars represent mean ± SEM.

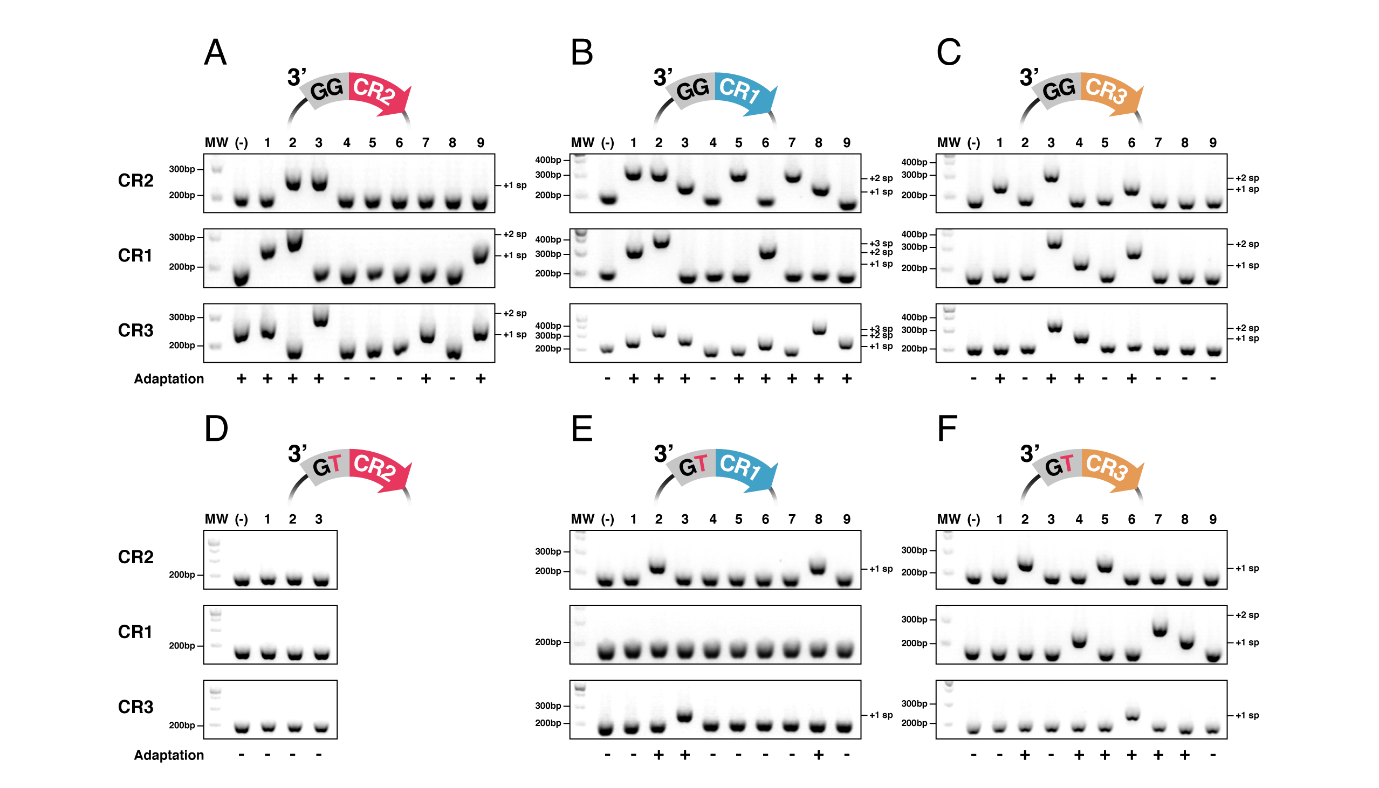

**Figure S6. All three CRISPR arrays of PA10145 can expand following CRISPR-mediated plasmid loss.** (A-F) Each of the three CRISPR arrays was screened for expansion where the lowest band at 200 bp shows the unexpanded CRISPR. As denoted by (-), PCR controls presumed negative for adaptation were sampled from a resistant colony on the S plate of the latest day with surviving colonies. Nine (9) random antibiotic-sensitive colonies were screened for CRISPR expansion among the biological triplicates from NS plates at day 5. Colonies with expansion in any CRISPR are positive for adaptation. Each spacer acquired increments the band by 60 bp, denoted as +1sp, +2sp, and +3sp. Shown are AGE profiles of sensitive colonies obtained from exposure to T plasmids containing PS recognized by (A) CR2, (B) CR1, and (C) CR3; and NT plasmids with PS derived from (D) CR2, (E) CR1, and (F) CR3. (A) Notably, the sampled negative control for the T-CR2 set-up contained an expanded CR3. A PCR screen revealed that the colony no longer contained the plasmid (data not shown) indicating that the colony had spontaneously gained resistance.

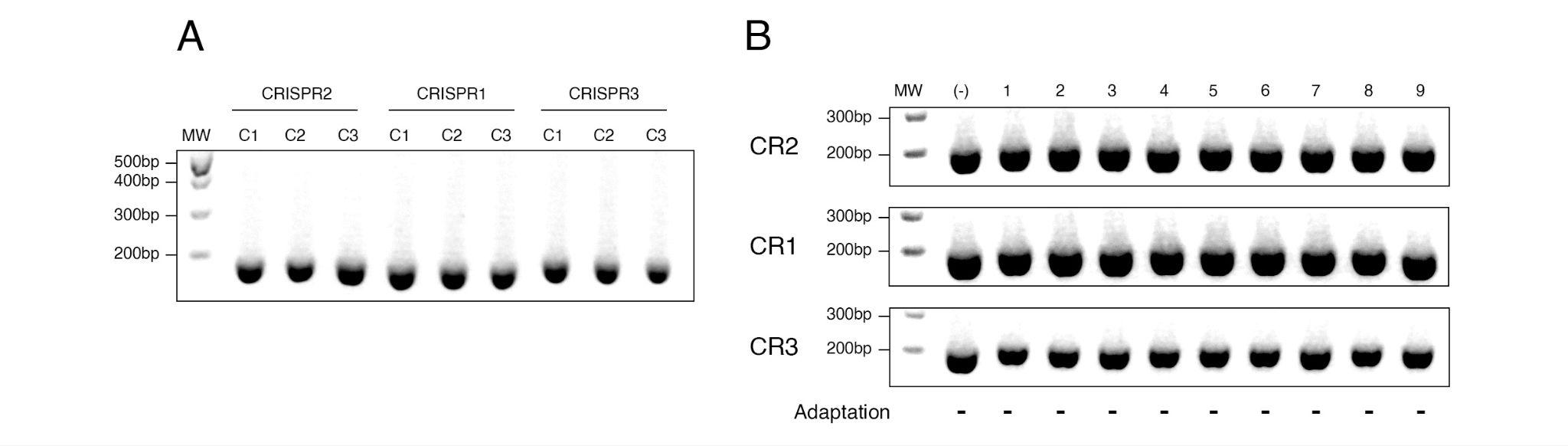

**Figure S7. No naïve adaptation was observed.** AGE profiles of CRISPR expansions for day 5 of the plasmid maintenance assay incubated with the control no PS vector using PCR templates obtained from (A) the bacterial population and (B) each antibiotic-sensitive colony.

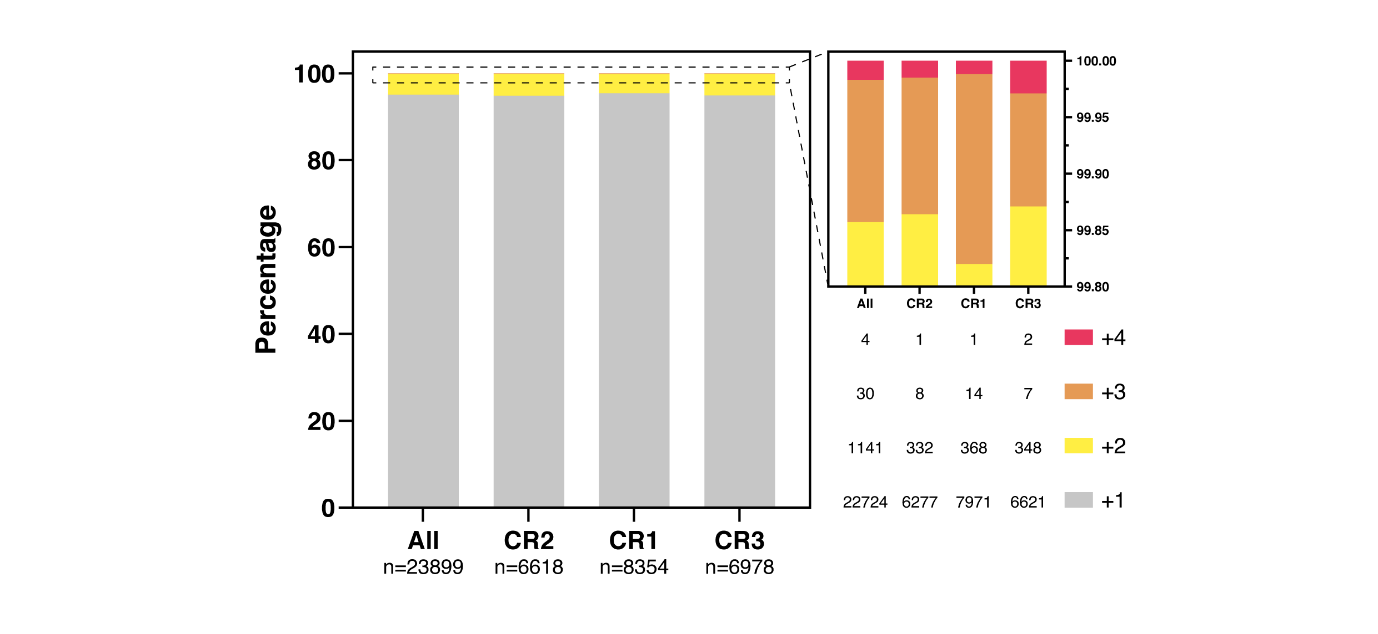

**Figure S8. Expanded arrays mostly acquire one spacer.** Sequencing statistics classified according to the number of newly acquired spacers in each expanded CRISPR array (n = 23,899). The frequency of these expansions is represented on the figure and enumerated on the lower right. Expanded arrays were binned to the three CRISPR arrays of PA10145 according to the two pre-existing spacers included in the sequenced amplicon.

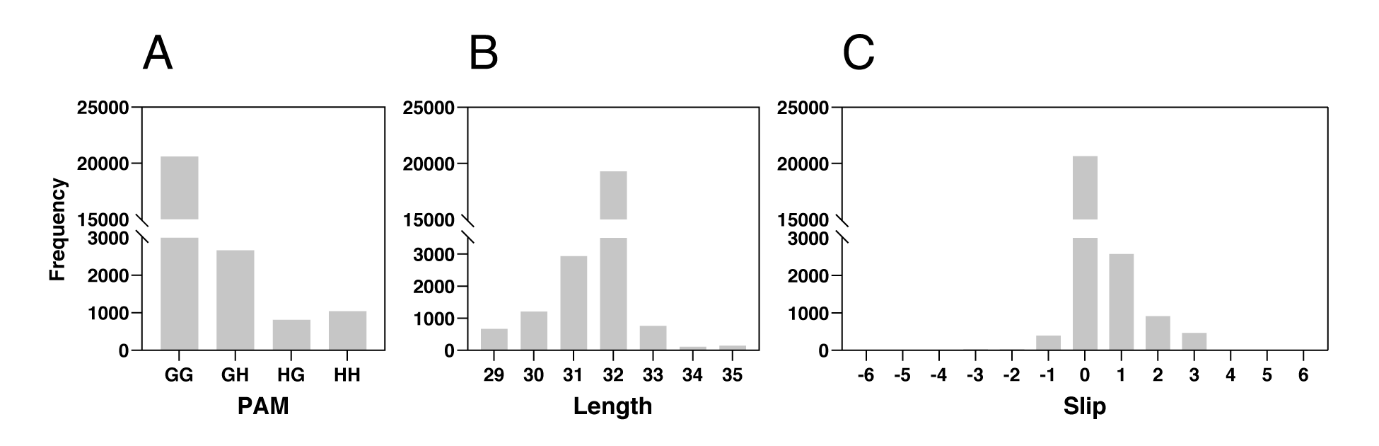

**Figure S9. Assessment of novel spacers.** (A) PAM frequencies of PS targeted by novel spacers. Most of the spacers mapped to a PS flanked by a 3’-GG PAM (82.02%). (B) Frequencies of spacers according to length. The majority (76.76%) exhibited the canonical length of 32 bp. Errors in sequencing might have introduced indels. (C) Slips are defined as the inaccurate shifting of the adaptation complex as it obtains a dsDNA substrate for incorporation into a CRISPR array [[40]](https://www.zotero.org/google-docs/?DL7loO). PAMs were corrected and assessed for slips with respect to probable correct PAMs 6 bp upstream and downstream. Corrected spacers now contain a canonical PAM (99.80%).

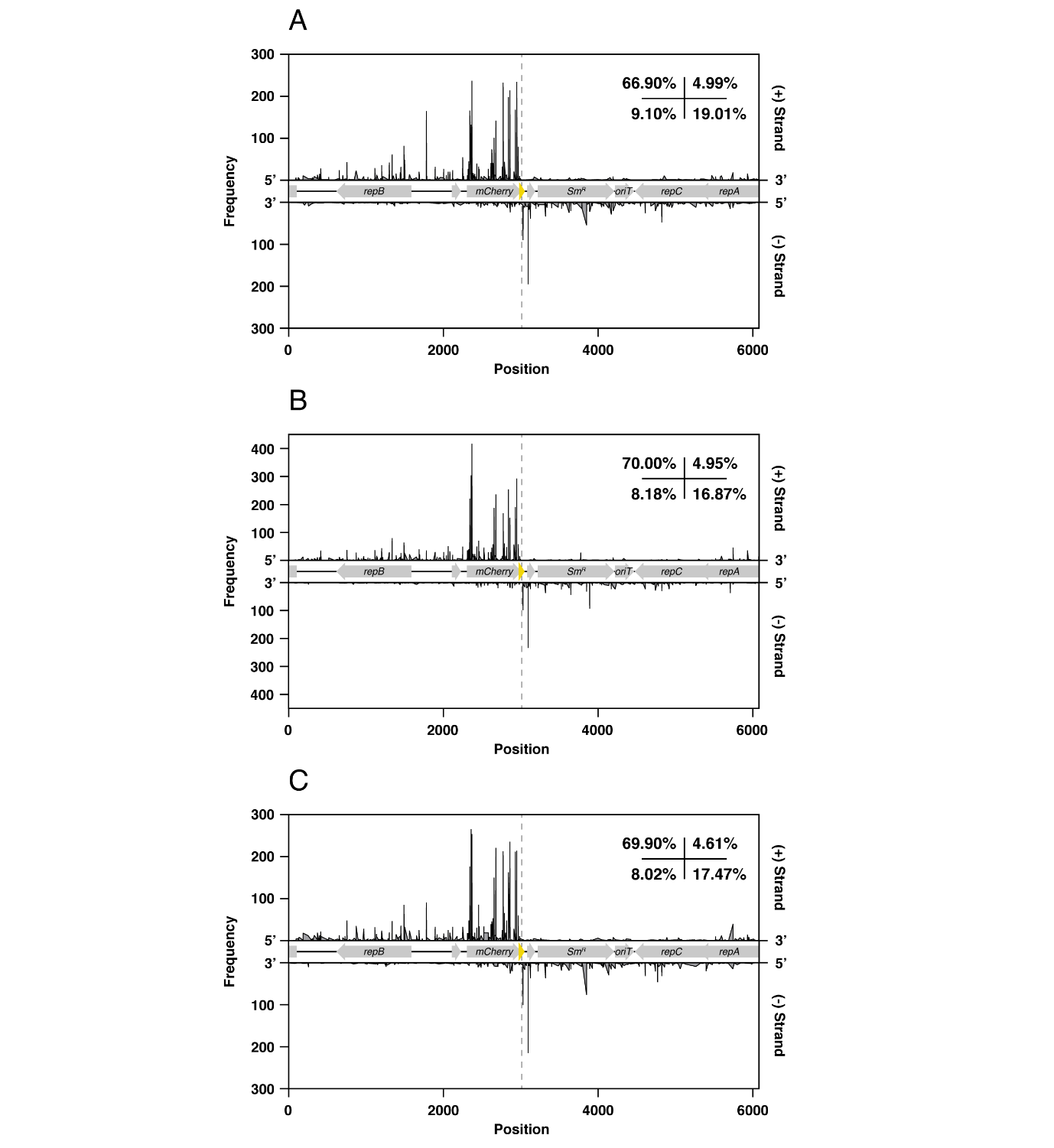

**Figure S10. The isolated CR3 participates in interference-mediated adaptation.** Mapping novel spacers acquired by (A) CR2, (B) CR1, and (C) CR3 to their protospacer counterparts in the T plasmid revealed a consistent trend in acquisitional frequency.

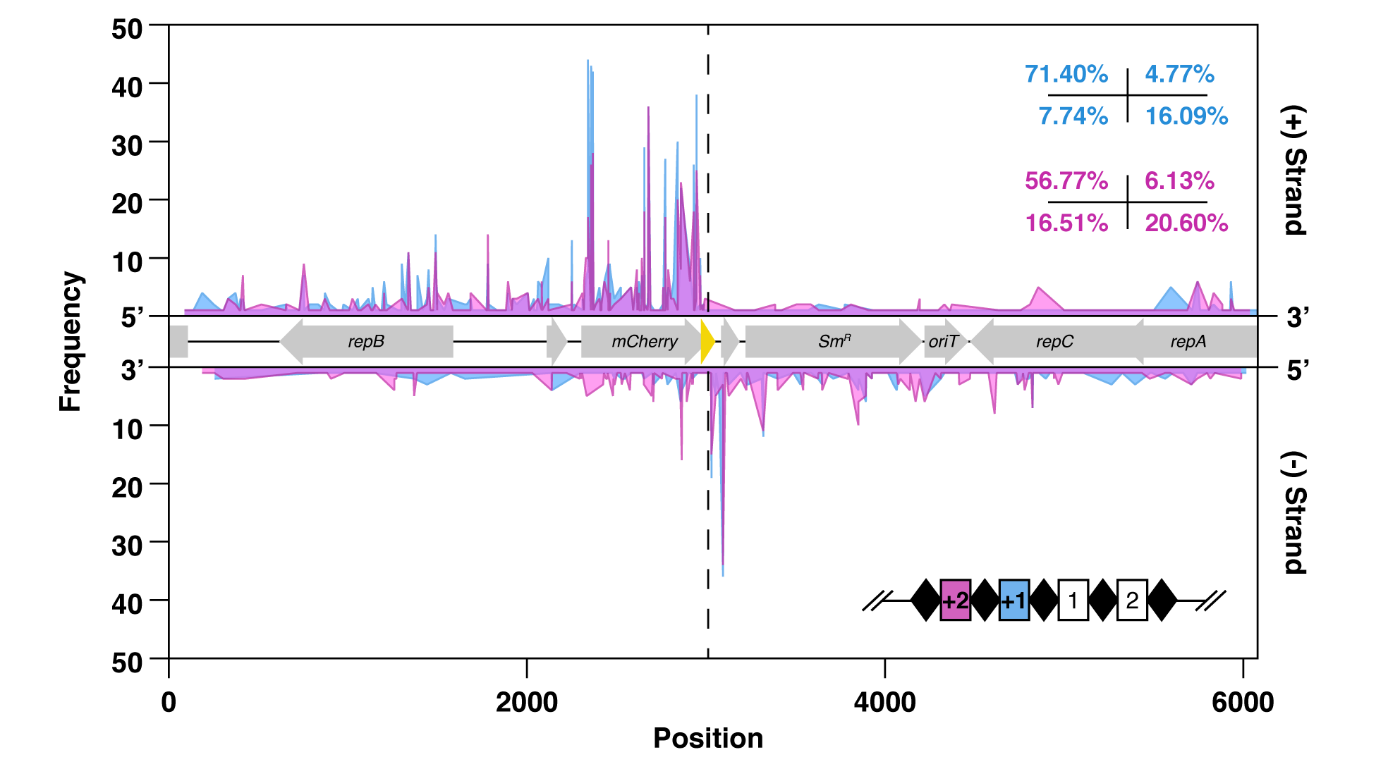

**Figure S11. Spacers are acquired sequentially during priming.** Protospacer distribution of the first (+1) and second (+2) spacers within the same expanded array with at least two newly-acquired spacers. A shift in distribution could be observed where higher percentages of +2 spacers are located in the target (-) strand and in the 3' direction of the target protospacer relative to the +1 spacer.

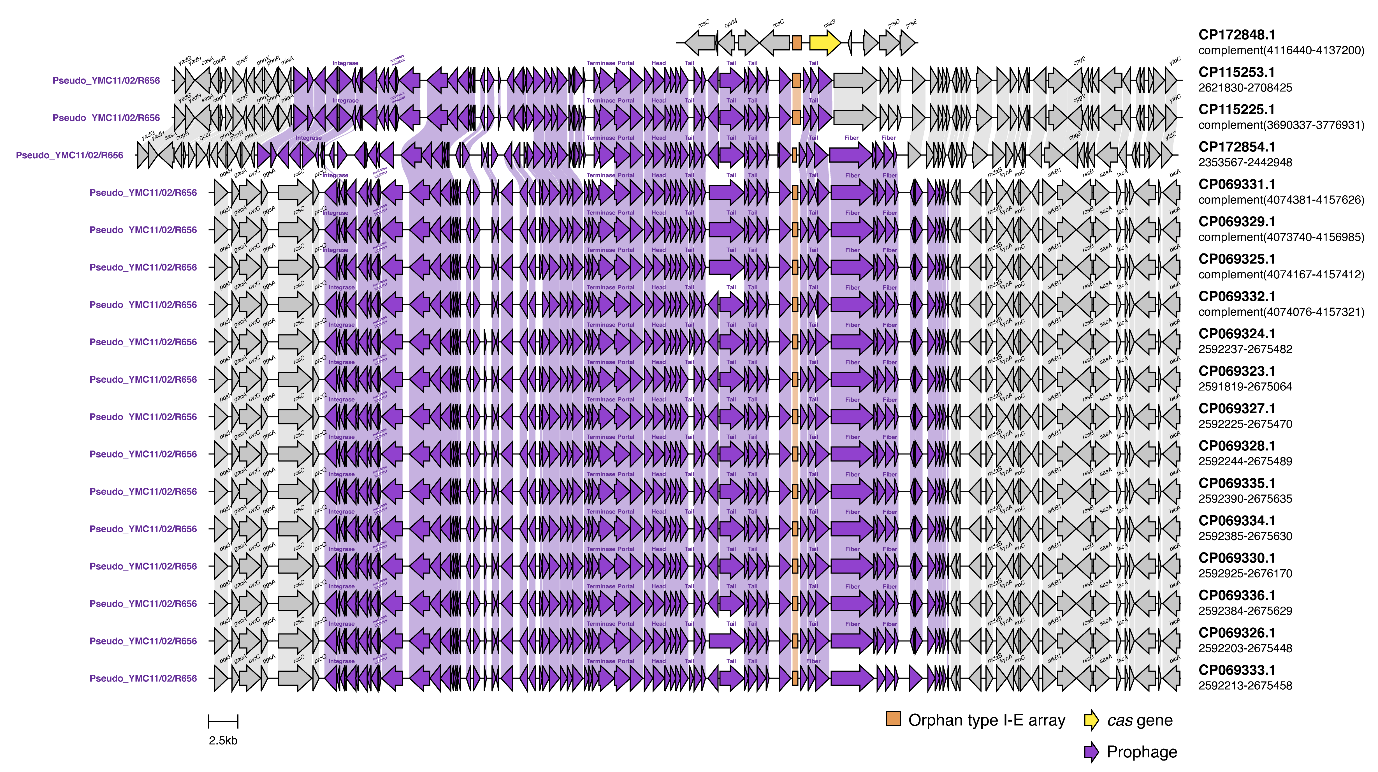

**Figure S12. Neighborhood alignments of all orphan I-E CRISPRs**. The majority of orphan I-E CRISPR arrays are embedded within prophages, likely contributing to their mobility across *Psa* genomes. Prophages were annotated by PHASTEST [[55]](https://www.zotero.org/google-docs/?7Ednou) with the closest identified prophage match indicated for each region. At the top, one other orphan I-E array emerged from reductive evolution of *cas*.

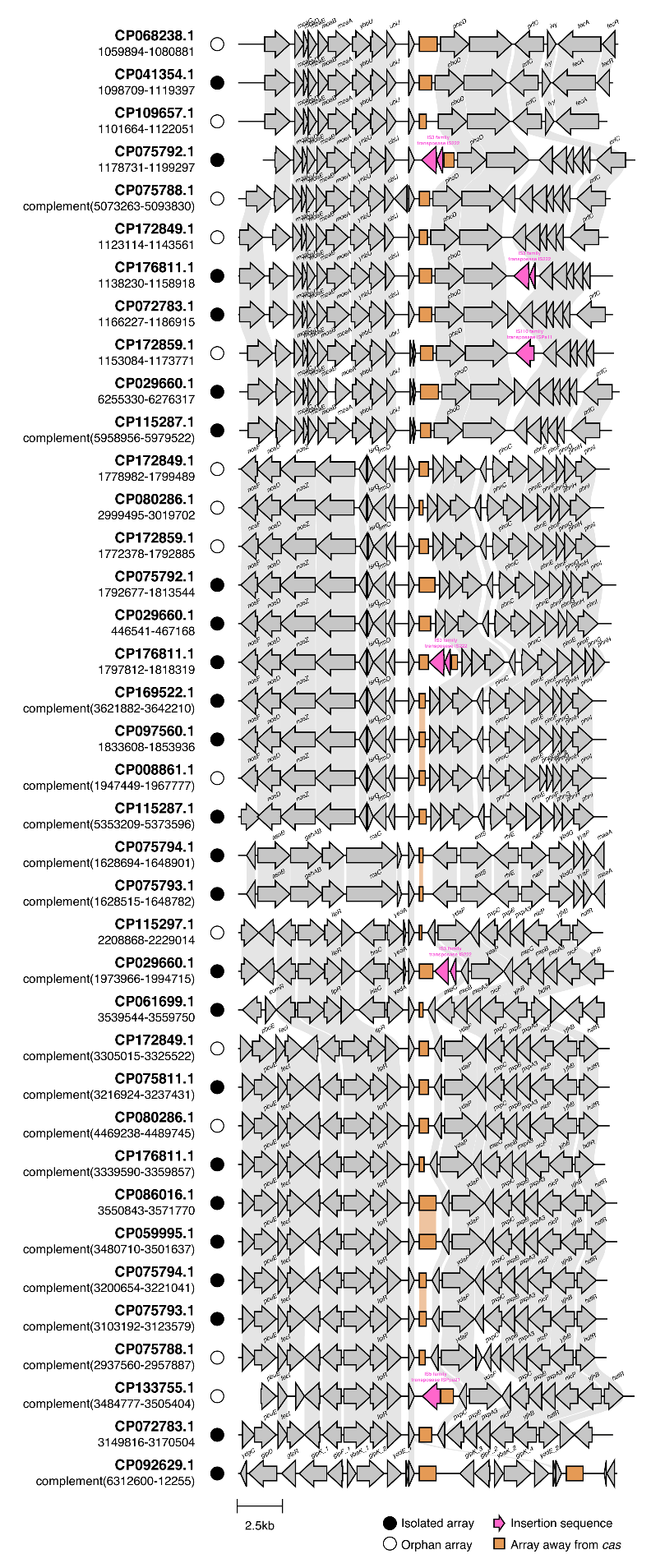

**Figure S13. Orphan and isolated I-F arrays located in sites apart from *ybaK-mmsR*.** Some isolated (n = 24) and orphan (n = 14) CRISPR arrays are found at distinct locations from PA10145-CR3. These arrays also have leader sequences that are >70% similar to the *cas*-adjacent CR2 leader. Two seemingly duplicated arrays can be found at the circularization junction of CP092629.1 (bottommost), possibly due to erroneous genome assembly.

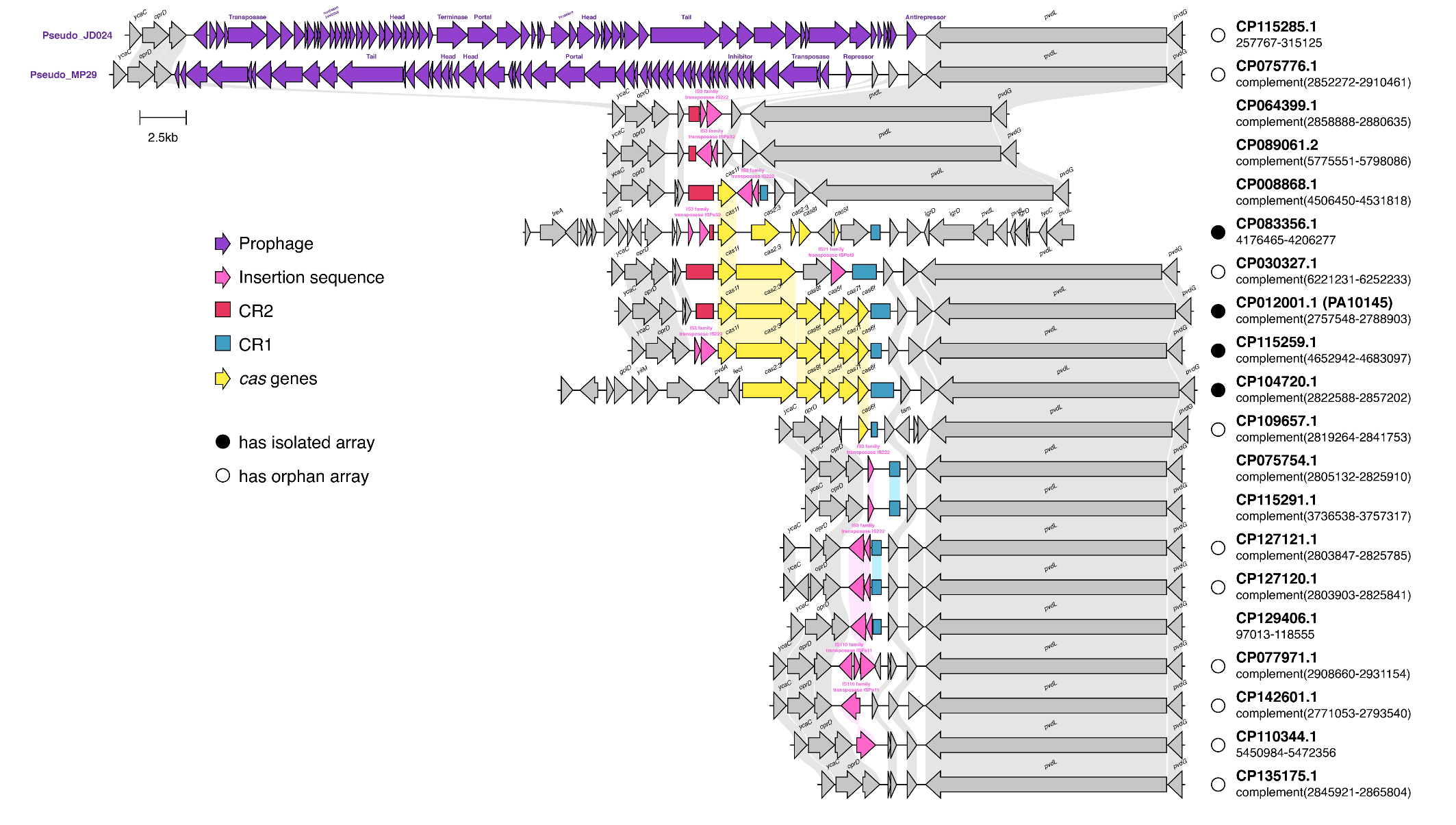

**Figure S14.** **Orphan arrays can emerge from *cas* reductive evolution.** The conserved *oprD-pvdL* region containing the type I-F CRISPR-Cas system was assessed across *Psa* genomes. Multiple *cas* genes were found to be deleted, leading to the emergence of orphaned CR2 and CR1. These may have also resulted in isolated I-F arrays to become orphaned, indicated as empty circles beside each genome accession.

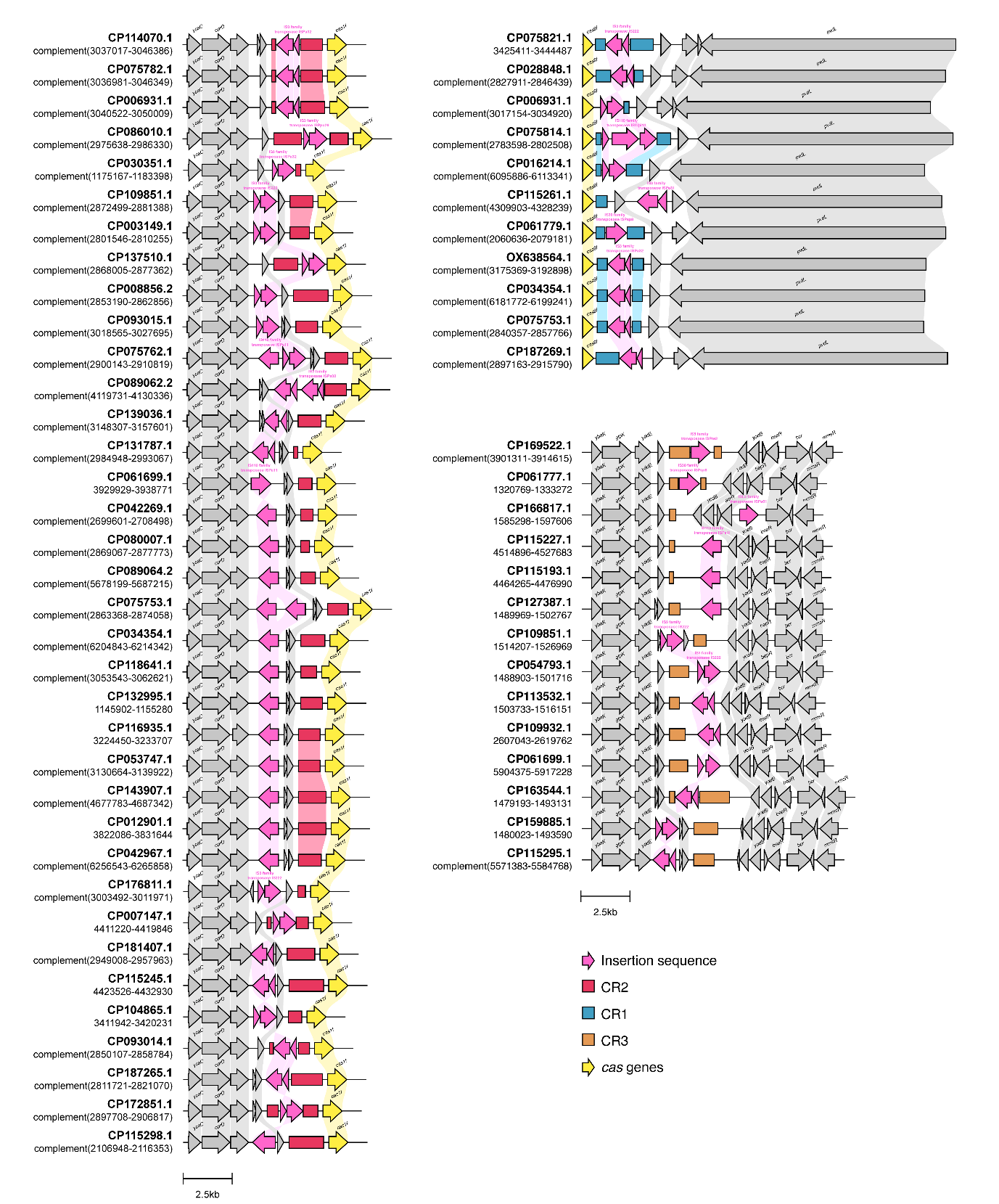

**Figure S15. Insertion sequences can be found in the CR2, CR1, and CR3 neighborhoods.** Insertion sequences (IS) were detected through Prokka [[54]](https://www.zotero.org/google-docs/?Un53OR) annotation. IS were found to be adjacent to I-F CRISPR arrays, or embedded within the leader or the array itself which leads to CRISPR disruption.

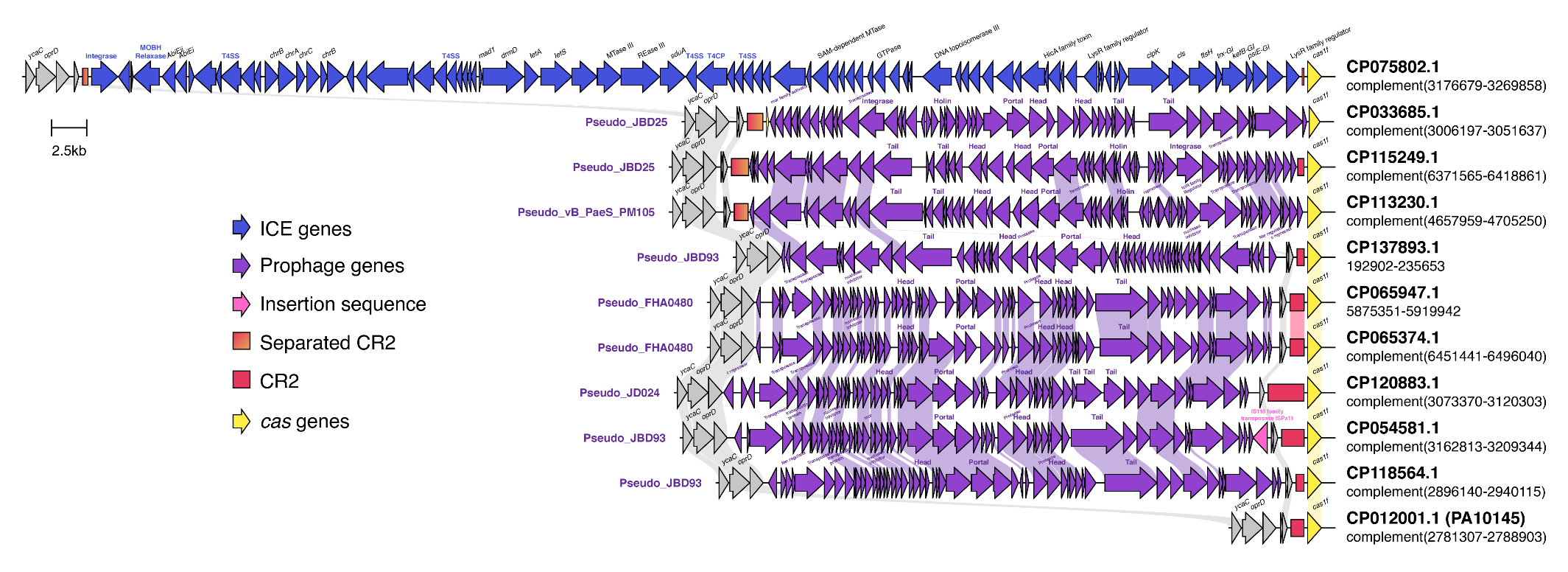

**Figure S16. Large MGE insertions in the CR2 neighborhood.** Integrative conjugative elements (ICE) and prophages were found within *oprD-cas1f*. An ICE recognized by ICEberg [[56]](https://www.zotero.org/google-docs/?4HTCbF) was determined to contain an intact integrase, relaxase, and T4SS machinery. Prophages were annotated by PHASTEST [[55]](https://www.zotero.org/google-docs/?ymeIXO) with the closest prophage match indicated per region. Insertion sequences were identified via Prokka [[54]](https://www.zotero.org/google-docs/?QsZ4cB) annotation. Some of these large MGEs are inserted within the CR2 array or the intergenic region between CR2 and *cas1f*, resulting in an isolated array that had emerged from a separated CR2.

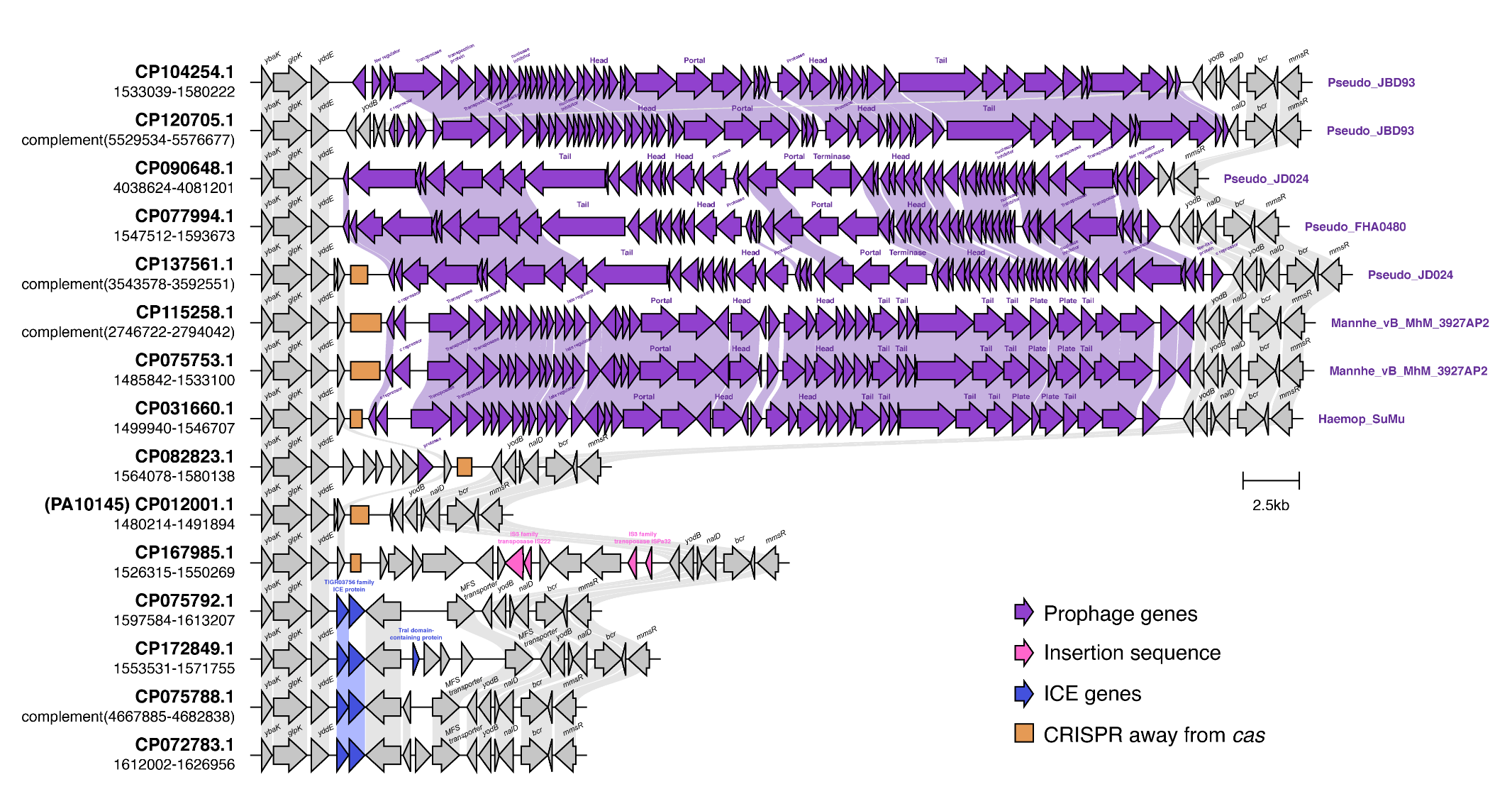

**Figure S17.** **Large MGE insertions in the CR3 neighborhood.** Various MGEs were found to be inserted within the *ybaK-mmsR* locus suggesting that insertion within this region can be tolerated by *Psa.* Such MGEs may also facilitate the mobility of CR3 across genomes. Intact prophages were detected by PHASTEST [[55]](https://www.zotero.org/google-docs/?6eVfnQ) with the closest prophage match indicated for each genomic region. Insertion sequences were identified via Prokka [[54]](https://www.zotero.org/google-docs/?hO5ugk) annotation. Annotated putative ICE genes were obtained from existing annotations of its genome.

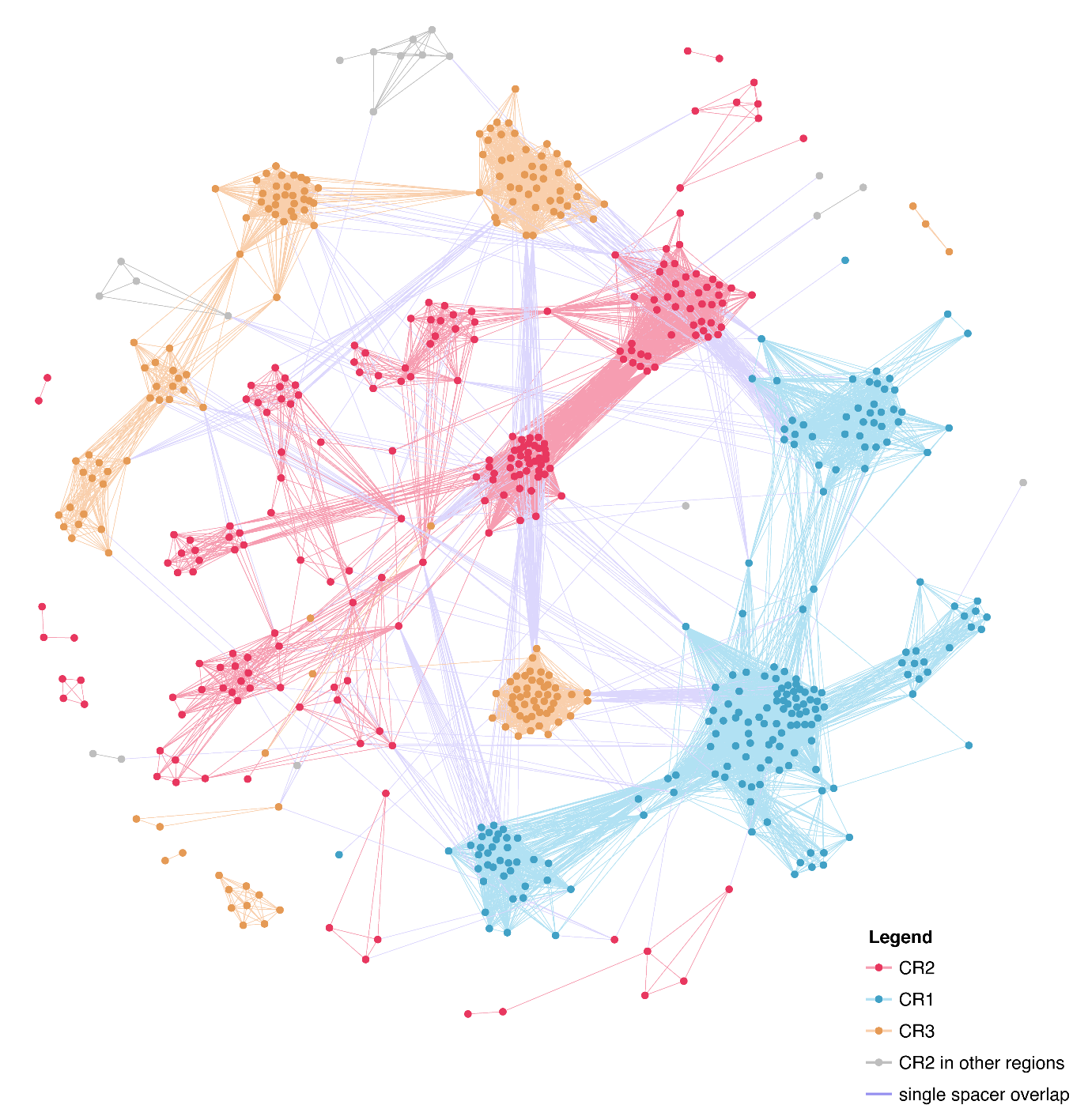

**Figure S18.** **Spacer sharing network of all *Psa* I-F CRISPR arrays**. Disrupted CRISPR arrays were excluded prior to network generation through the CCTK package [[42]](https://www.zotero.org/google-docs/?XVl9NK). Each node represents an array with a unique spacer profile. Node proximity is dictated by Jaccard similarity of spacers between two CRISPR arrays. Nodes within each CRISPR subset mostly cluster together yet minimally overlap with other subsets as shown by inter-subset edges that represent only a single shared spacer.

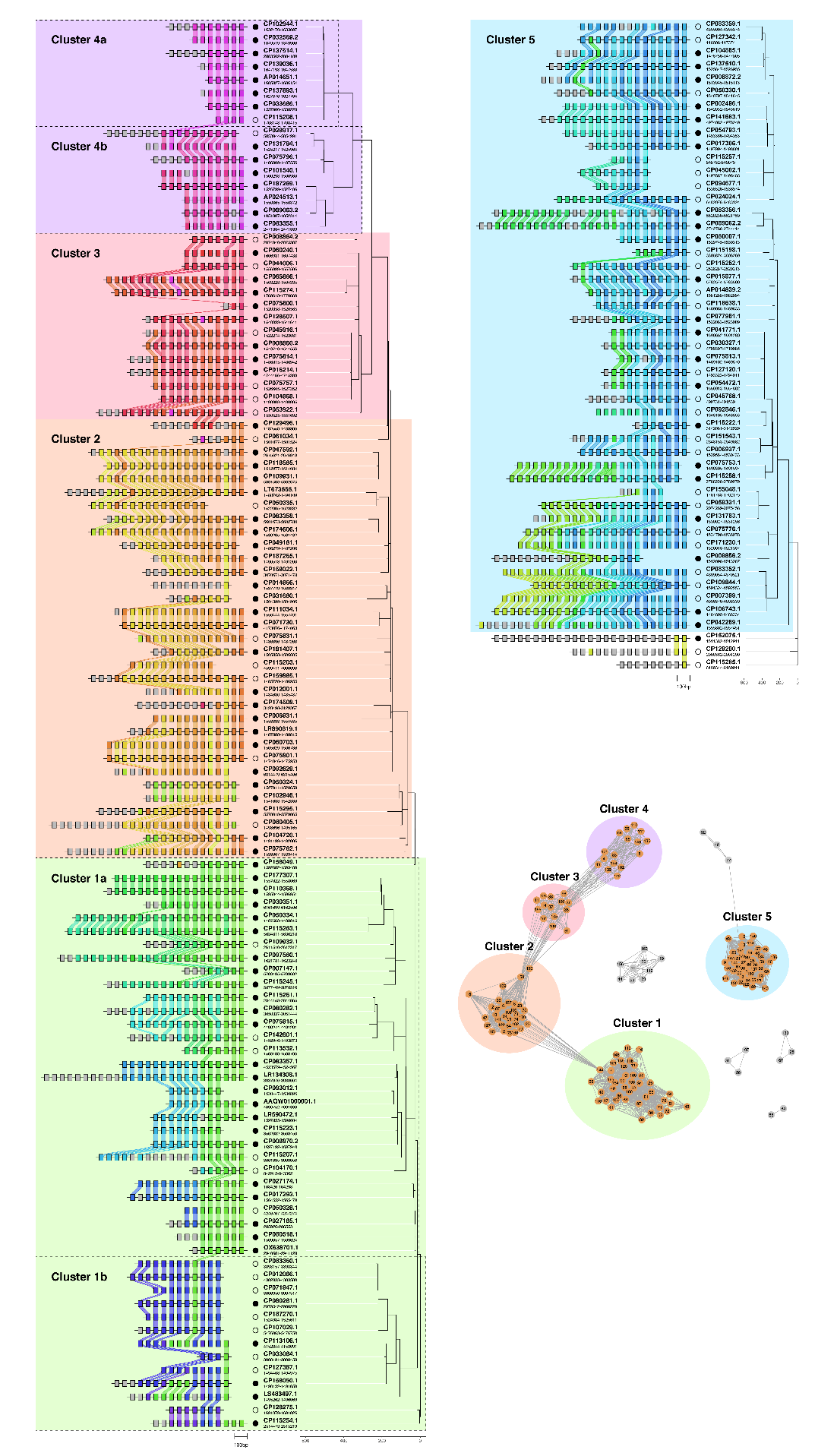

**Figure S19.** **CR3 maximum parsimony tree and spacer plots.** Clusters from the CR3 spacer sharing network with the fivemost number of arrays were subjected to CRISPRtree generation [[42]](https://www.zotero.org/google-docs/?7xtdTh). The quaternary clustering of the sub-network with the most number of arrays (Clusters 1-4) coincided with the generated maximum parsimony tree and spacer alignments. These clusters were used to classify the unique CR3 lineages harbored by each *Psa* genome.

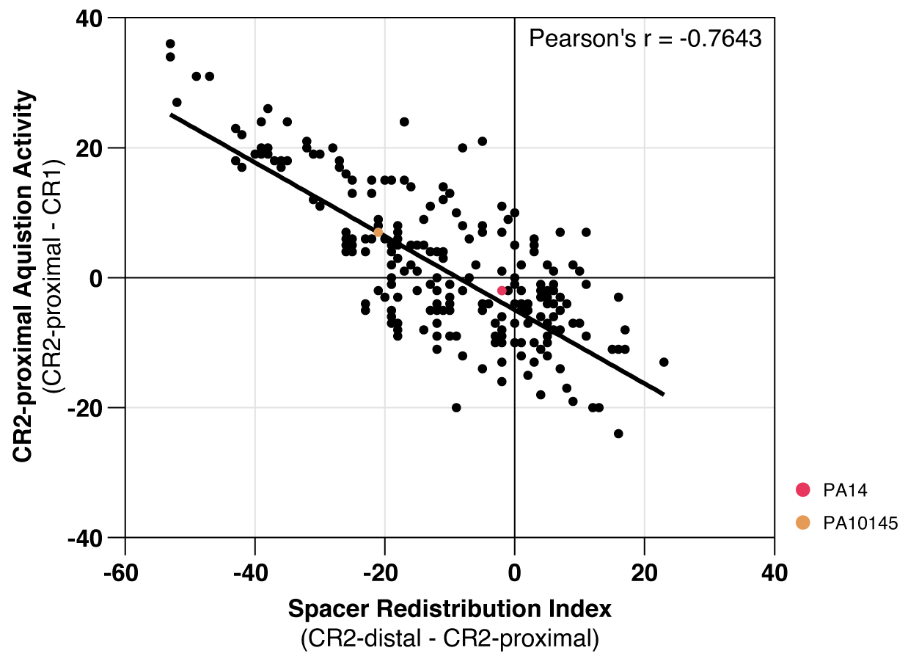

**Figure S20. Spacer acquisition at proximal CR2 arrays is redirected to distal CR2.** For each *P. aeruginosa* genome, the acquisition activity of CR2-proximal was plotted against its spacer redistribution index. Acquisition activity of CR2-proximal was defined as the difference in the number of spacers between CR2-proximal and CR1. Spacer redistribution index is the degree to which the presence of a CR2-distal has redirected spacer acquisition from CR2-proximal. Such a measure was designated to be the difference in the number of spacers between CR2-distal and CR2-proximal. Should multiple CR2-distal arrays exist in the genome, the array with the highest number of spacers was used. Genomes with disrupted or orphan CRISPR arrays were excluded, resulting in a plot of 393 genomes. Pearson's r revealed a strong negative correlation (r = -0.7643) between the two variables. PA14 and PA10145 are plotted as the red and orange dots, respectively
